# Neuritin-1 is an adipokine that sustains insulin sensitivity through adipocyte FGFR1–PPARγ signaling

**DOI:** 10.64898/2026.11.20.768237

**Authors:** Ikjun Lee, Dongjoo You, Louise Lantier, Owen P McGuinness, Byung Chul Jung, Diana L Alba, Suneil K Koliwad, Sona Kang

## Abstract

Obesity remodels the adipose secretome, yet the adipocyte-derived signals that preserve insulin sensitivity remain incompletely defined. Here we identify neuritin (NRN1), a GPI-anchored protein, as an obesity-suppressed adipokine that protects against insulin resistance. In white adipose tissue, *Nrn1* expression increases after adipocyte-specific deletion of DNA methyltransferase 3A (DNMT3A), which occupies the *Nrn1* promoter. In obese mice, higher adipose DNMT3A and *Nrn1* promoter methylation accompany lower *Nrn1* expression, reduced NRN1 release from white adipose tissue and lower circulating NRN1. In humans, adipocyte *NRN1* expression and plasma NRN1 concentrations are inversely associated with body mass index. Increasing NRN1 expression in white adipose tissue or raising circulating NRN1 improves insulin sensitivity and glucose homeostasis in obese mice without altering adiposity, whereas adipocyte-specific *Nrn1* deletion exacerbates diet-induced insulin resistance. NRN1 enhances adipose insulin signaling and glucose uptake, raises circulating adiponectin, and restrains inflammation, adipocyte hypertrophy, and lipolysis. Mechanistically, NRN1 engages FGFR1 on adipocytes, stimulating receptor phosphorylation and downstream signaling. For cell-expressed NRN1, this association depends on native GPI-anchor processing. Pharmacological or genetic FGFR1 inhibition abrogates NRN1-induced insulin sensitization *in vitro*, and adipocyte-specific *Fgfr1* deletion largely abolishes NRN1’s adipose and systemic effects *in vivo*. This pathway converges on PPARγ: PPARγ antagonism blocks NRN1-enhanced insulin-stimulated glucose uptake, whereas rosiglitazone restores this response in *Nrn1*-deficient adipocytes and largely corrects insulin resistance in knockout mice. Thus, NRN1 is a previously unrecognized adipokine that acts as a non-FGF signal through adipocyte FGFR1 and PPARγ to preserve adipose function and systemic insulin sensitivity.

## Introduction

Insulin resistance is central to type 2 diabetes (T2D) and contributes to cardiovascular disease and metabolic dysfunction-associated steatotic liver disease (MASLD)^1-3^. Although obesity is a major driver, fat mass alone does not determine insulin sensitivity: some people with obesity retain relatively preserved metabolic health^4^, and adipose-targeted manipulations in mice improve insulin action despite increased or unchanged adiposity^5-8^. Together, these findings highlight adipose tissue function as an important, modifiable contributor to systemic insulin sensitivity. This distinction has clinical relevance because some current therapies cause weight gain or other adverse effects^9-11^, while weight loss may not be appropriate for every patient, particularly when preserving lean mass and physical function is a priority^1^. Identifying endogenous pathways that improve insulin responsiveness without reducing adiposity could therefore provide a complementary therapeutic strategy.

Adipose tissue regulates systemic insulin sensitivity not only by buffering excess lipid but also by secreting hormones, peptides and lipid mediators that communicate with the liver, skeletal muscle, pancreas and brain^12-19^. The discovery of leptin established adipocytes as endocrine cells^20, 21^, and adiponectin and many other adipokines have since been implicated in systemic metabolic control^17, 22-25^. Obesity profoundly remodels the adipocyte secretome: circulating leptin rises, whereas adiponectin and adipsin fall, and loss of adiponectin is causally linked to insulin resistance^16, 26-30^. However, whether additional adipocyte-derived factors are diminished in obesity, and whether their loss contributes to insulin resistance, remains largely unexplored.

Glycosylphosphatidylinositol-anchored proteins (GPI-APs) regulate receptor signaling, cell adhesion, immune function and neuronal development^31^. Regulated cleavage can release GPI-APs from the membrane, enabling local and distant actions^31-33^. Neuritin-1 (NRN1) is a small GPI-AP best known for its roles in the nervous system, where it contributes to synaptic plasticity, neurite outgrowth, and neuronal survival^34-36^. Less is known about its functions outside the nervous system. Human genetic studies have linked the *NRN1* locus to body mass index (BMI) and cardiometabolic stroke^37^, suggesting a possible metabolic role for NRN1. We previously showed that adipocyte-specific deletion of the *Dnmt3a* protects mice from diet-induced insulin resistance^8^. Transcriptomic profiling of epididymal white adipose tissue (eWAT) from these mice identified *Nrn1* as one of the most strongly upregulated genes. These observations prompted us to ask whether NRN1 links DNMT3A-dependent gene regulation to insulin responsiveness in adipose tissue.

GPI-APs lack transmembrane and cytoplasmic domains and can relay signals through partners such as receptor tyrosine kinases^38-40^. The receptor through which NRN1 acts on adipocytes, however, is unknown. Using phosphor-kinase array screening, we identified fibroblast growth factor receptor 1 (FGFR1), a central regulator of adipocyte metabolic homeostasis^41-45^. In adipose tissue, FGFR1c mediates the insulin-sensitizing actions of FGF21, with β-Klotho as co-receptor, and of FGF1, with heparan sulfate proteoglycans^46-48^. Both pathways intersect with PPARγ^45, 49-51^, a master regulator of adipogenesis, insulin responsiveness, lipid storage and adipocyte endocrine function. We therefore asked whether NRN1, an endogenous non-FGF protein, engages adipocyte FGFR1–PPARγ signaling to preserve adipose function and systemic insulin sensitivity.

Here we identify NRN1 as an adipokine that is suppressed in obesity and protects against insulin resistance. DNMT3A occupies the *Nrn1* promoter. In obese mice, increased methylation of this promoter in eWAT coincides with reduced adipose *Nrn1* expression and lower circulating NRN1. In humans, adipocyte *NRN1* expression and circulating NRN1 concentrations are each inversely associated with BMI. Restoring NRN1 improves insulin sensitivity in obese mice, whereas adipocyte-specific deletion worsens it, in both cases without altering adiposity. Mechanistically, NRN1 engages FGFR1 on adipocytes, a non-FGF input into this pathway, and activates a PPARγ-dependent program that enhances glucose uptake and insulin signaling while limiting inflammation and lipolysis. These findings define a DNMT3A–NRN1–FGFR1– PPARγ axis through which obesity erodes adipose insulin responsiveness, and establish NRN1 as a protective adipokine.

## Results

### Obesity suppresses adipocyte NRN1 expression and release in association with DNMT3A-linked promoter methylation

Adipocyte-specific *Dnmt3a* knockout (*Dnmt3a*ᴬᴷᴼ) mice show enhanced insulin sensitivity on a high-fat diet (HFD)^8^. To identify genes that might mediate this effect, we profiled gene expression in eWAT from knockout and control mice fed a HFD. *Nrn1* was among the most strongly upregulated genes. In adipocytes isolated from knockout eWAT, *Nrn1* mRNA and NRN1 protein were approximately fivefold and threefold higher, respectively, than in controls (**Figs. 1A,B**). Reciprocal perturbations in cultured adipocytes were consistent with DNMT3A-mediated repression of *Nrn1*, as *Dnmt3a* knockdown increased *Nrn1* expression by approximately 1.7 times the control level, whereas DNMT3A overexpression reduced it by approximately 50% **(Figs. 1C,D)**.

**Figure 1.**
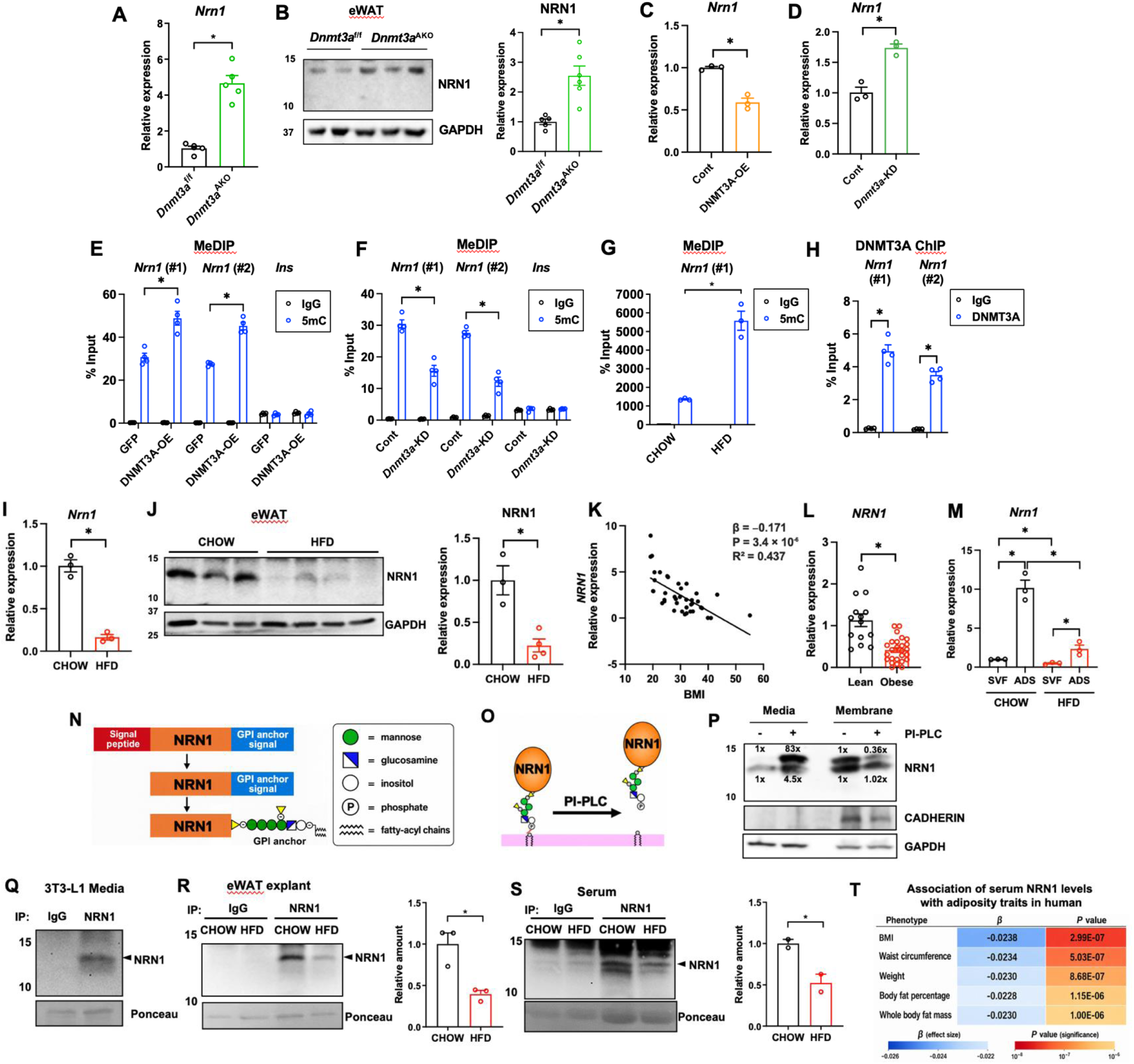
Obesity suppresses the expression and secretion of NRN1, an adipocyte-enriched GPI-anchored protein. **(A)** *Nrn1* mRNA levels in isolated from epididymal white adipose tissue (eWAT) of HFD-fed *Dnmt3a*^f/f^ and adipocyte-specific *Dnmt3a*-knockout (*Dnmt3a*^AKO^) mice (*n* = 5 per group). **(B)** NRN1 protein levels in eWAT from *Dnmt3a*^f/f^ (*n* = 5) and *Dnmt3a*^AKO^ (*n* = 6) mice. **(C, D)** *Nrn1* mRNA levels in 3T3-L1 adipocytes transduced with lentiviruses for *Dnmt3a* knockdown (KD**) (C)** or DNMT3A overexpression (OE) **(D)** (*n* = 3). 5-Methylcytosine (5mC) enrichment at two regions of the *Nrn1* promoter (regions 1 and 2) and the *Ins1* locus (negative control), measured by methylated DNA immunoprecipitation followed by quantitative PCR (MeDIP–qPCR), in 3T3-L1 adipocytes with DNMT3A-OE **(E)** or *Dnmt3a*-KD **(F)** (*n* = 4). **(G)** 5mC enrichment at region 1 of the *Nrn1* promoter in eWAT from chow- and HFD-fed mice (*n* = 3 per group). **(H)** DNMT3A occupancy at the *Nrn1* promoter in 3T3-L1 adipocytes, assessed by DNMT3A ChIP-qPCR (*n* = 4). *Nrn1* mRNA levels (**I**; *n* = 3 per group) and representative immunoblots and quantification of NRN1 protein (**J**; chow, *n* = 3; HFD, *n* = 4) in eWAT from chow- and HFD-fed mice. **(K)** Correlation between *NRN1* mRNA levels in adipocytes isolated from human WAT and body mass index (BMI; *n* = 40 individuals). The association between BMI and relative NRN1 expression in isolated human adipocytes was analyzed using ordinary least-squares linear regression. The unstandardized regression coefficient (β), 95% confidence interval (CI), two-sided P value, and R² are reported. **(L)** *NRN1* mRNA levels in WAT from lean (BMI <25; n =14) and obese individuals (BMI ≥25, n = 26). **(M)** *Nrn1* mRNA levels in mature adipocytes and the stromal vascular fraction (SVF) isolated from eWAT of chow- and HFD-fed mice (*n* = 3 per group). **(N)** Schematic illustrating NRN1 maturation and the composition of its GPI anchor. **(O)** Schematic illustrating phosphatidylinositol-specific phospholipase C (PI-PLC)-mediated release of NRN1 from the plasma membrane. **(P)** Immunoblot analysis of membrane-associated and released NRN1 in 3T3-L1 adipocytes treated with or without PI-PLC. **(Q)** Detection of soluble NRN1 in conditioned medium from 3T3-L1 adipocytes by immunoprecipitation. **(R)** Relative NRN1 abundance in conditioned medium from eWAT explants of chow- and HFD-fed mice. (*n* = 3 per group). **(S)** Relative NRN1 abundance in serum from chow- and HFD-fed mice (*n* = 3 per group). **(T)** Heatmap summarizing associations of the NRN1 abundance with anthropometric and adiposity-related traits. Data are presented as mean ± s.e.m. Pairwise comparisons were performed using two-tailed unpaired Student’s *t*-tests.

DNMT3A overexpression also increased methylation of the *Nrn1* promoter, whereas *Dnmt3a* knockdown reduced it (**Figs. 1E,F**), in line with the reciprocal changes in *Nrn1* expression. (**Figs. 1C,D**). In eWAT, where DNMT3A abundance increases in obesity^8^, the same promoter region was hypermethylated in obese mice relative to lean controls (**Fig. 1G**). Chromatin immunoprecipitation (ChIP) assay detected DNMT3A occupancy at the *Nrn1* promoter in adipocytes (**Fig. 1H**). These findings identify *Nrn1* as a likely direct target of DNMT3A and suggest that promoter methylation contributes to its gene repression in obesity.

NRN1 was also reduced in obesity. Both *Nrn1* transcript and NRN1 protein levels were approximately 80% lower in eWAT from diet-induced obese and *ob/ob* mice than in their respective lean controls (**Figs. 1I,1J and Supplementary Fig. 1A**). A similar relationship was evident in humans. In a cohort^52^ with body mass index (BMI) values ranging from 18.5 to 53 kg m⁻², *NRN1* mRNA levels displayed a strong inverse association with BMI (β = −0.171, *P* = 3.4 × 10⁻⁶, **Fig. 1K**). Adipocyte *NRN1* expression was significantly lower in individuals with overweight or obesity (BMI ≥25 kg m⁻²) than in lean individuals (**Fig. 1L**). Fractionation of mouse eWAT showed that *Nrn1* is enriched in adipocytes relative to the stromal vascular fraction in both lean and obese mice. The obesity-associated reduction in expression was more pronounced in adipocytes (**Fig. 1M**). *Nrn1/NRN1* mRNA expression also increased during differentiation of mouse 3T3-L1 and human Simpson–Golabi–Behmel syndrome (SGBS) preadipocytes (**Supplementary Figs. 1B,C**), consistent with enrichment in the adipocyte fraction.

We next asked whether adipocytes release NRN1. During maturation in the endoplasmic reticulum, NRN1 loses its N-terminal signal peptide and acquires a GPI anchor in place of its C-terminal anchoring signal (**Fig. 1N**). This anchor tethers mature NRN1 to the outer leaflet of the plasma membrane^34, 53^ (**Fig. 1N**). Treating mature 3T3-L1 adipocytes with bacterial phosphatidylinositol-specific phospholipase C (PI-PLC), which cleaves GPI anchors^54^, reduced membrane-associated NRN1 and increased NRN1 in the conditioned medium (**Figs. 1O,1P**). We also detected NRN1 in conditioned medium from untreated adipocytes, indicating that adipocytes release NRN1 under basal conditions.

NRN1 release was also reduced in obesity: its level in conditioned medium from obese eWAT explants were approximately 60% lower than that from lean controls (**Fig. 1R**), and circulating NRN1 was approximately 50% lower in obese mice (**Fig. 1S**). In humans, plasma NRN1 abundance in a published proteomic dataset was inversely associated with BMI and several other measures of adiposity^55^ (**Fig. 1T**). Together, these findings identify adipocytes as a source of extracellular NRN1 and show that lower adipose NRN1 expression in obesity is accompanied by lower circulating NRN1.

### NRN1 enhances adipocyte insulin sensitivity in a GPI-anchor-dependent manner

We first tested whether extracellular NRN1 enhances the response of adipocytes to insulin. In mature 3T3-L1 adipocytes, recombinant NRN1 (reNRN1) enhanced insulin-stimulated 2-deoxyglucose uptake in a dose-dependent manner, both under control conditions and in the presence of TNFα (TNF), which induces inflammation-associated insulin resistance^56^ (**Fig. 2A**). Conditioned medium from NRN1-overexpressing cells also enhanced insulin-stimulated glucose uptake in recipient adipocytes (**Fig. 2B**). Together with the detection of NRN1 in conditioned medium from untreated adipocytes (**Fig. 1Q**), these findings support a autocrine action of adipocyte-derived NRN1.

**Figure 2.**
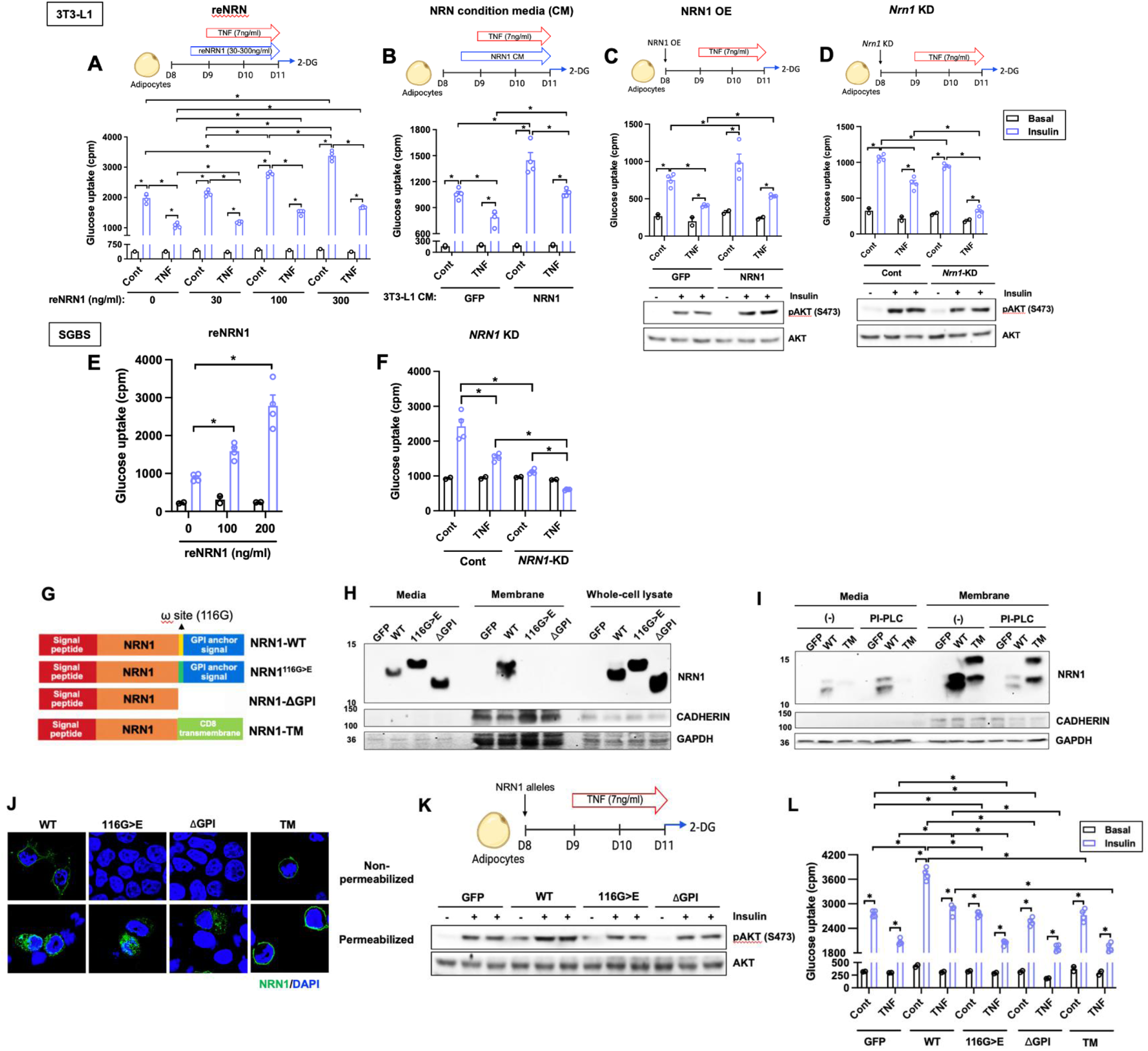
The insulin-sensitizing activity of adipocyte-expressed NRN1 depends on its native GPI-anchoring signal. **(A)** Glucose uptake of 3T3-L1 treated with various reNRN1 concentration, with or without TNFα (TNF; *n* = 2 for basal and *n* = 4 for insulin-stimulated conditions). **(B)** Glucose uptake in wild-type 3T3-L1 adipocytes treated with conditioned media collected from HEK293 cells from NRN1 or GFP, with or without TNF (*n* = 2 and 4, respectively). **(C) Top:** Glucose uptake in 3T3-L1 adipocytes overexpressing GFP or NRN1, with or without TNF treatment (*n* = 2 and 4, respectively). **Bottom:** AKT phosphorylation in 3T3-L1 adipocytes overexpressing GFP or NRN1 under basal or insulin-stimulated conditions (*n* = 1 and 2, respectively). **(D) Top:** Glucose uptake in control and *Nrn1*-knockdown 3T3-L1 adipocytes, with or without TNF treatment (*n* = 2 and 4, respectively). **Bottom:** AKT phosphorylation in control and *Nrn1*-knockdown 3T3-L1 adipocytes under basal or insulin-stimulated conditions (*n* = 1 and 2, respectively). **(E)** Glucose uptake of SGBS treated with various reNRN1 concentration, with or without TNF (*n* = 2 and 4, respectively). **(F)** Glucose uptake in control or *NRN1-*knockdown SGBS adipocytes, with or without TNF (*n* = 2 and 4, respectively). **(G)** Schematic domain structures of wild-type NRN1 (NRN1-WT) and the GPI-anchor-modified variants (NRN1^G116<E^, NRN1-ΔGPI, and NRN1-TM). **(H)** Immunoblot analysis of NRN1-WT, NRN1^G116<E^, and NRN1-ΔGPI in conditioned medium, membrane fractions and whole-cell lysates from HEK293T cells. **(I)** Immunoblot analysis of NRN1-WT and NRN1-TM in conditioned medium and membrane fractions from HEK293T cells treated with or without PI-PLC. **(J)** Immunocytochemical localization of NRN1-WT and the anchor-modified-variants at the plasma membrane under non-permeabilized conditions and intracellularly under permeabilized conditions. **(K)** AKT phosphorylation in 3T3-L1 adipocytes expressing GFP, NRN1-WT, NRN1^G116<E^, or NRN1-ΔGPI under basal and insulin-stimulated conditions (*n* = 1 for basal conditions and *n* = 2 for insulin-stimulated conditions). **(L)** Glucose uptake in 3T3-L1 adipocytes overexpressing GFP, NRN1-WT or the GPI-anchor-modified variants, with or without TNF, under basal or insulin-stimulated conditions (*n* = 2 and 4, respectively). Data are presented as mean ± s.e.m. Pairwise comparisons were performed using two-tailed unpaired Student’s *t*-tests.

We next examined how increasing or reducing NRN1 expression in adipocytes affected their response to insulin. An approximately fourfold increase in NRN1 (**Supplementary Fig. 2A**), within the range of the lean-versus-obese difference (**Figs. 1I,J**), enhanced the effects of insulin on both glucose uptake and AKT phosphorylation in mature 3T3-L1 adipocytes (**Fig. 2C**). Conversely, CRISPR–Cas9-mediated *Nrn1* knockdown impaired both responses in the presence and absence of TNF (**Fig. 2D**). Gain- and loss-of-function experiments in human SGBS adipocytes yielded concordant results (**Figs. 2E,F, and Supplementary Fig. 2B**), indicating that NRN1’s insulin-sensitizing effect is conserved between mouse and human adipocytes.

To test how GPI-anchor processing affects NRN1 localization and whether cell-surface targeting alone is sufficient to preserve its insulin-sensitizing activity, we compared wild-type NRN1 with three variants (**Fig. 2G**). NRN1-G116E carries a glycine-to-glutamate substitution at the predicted GPI attachment site (Gly116, the ω-site), a change expected to disrupt GPI-anchor attachment by transamidation^57-59^. NRN1-ΔGPI lacks the entire C-terminal GPI-anchoring signal. In NRN1-TM, we replaced the GPI-anchoring signal with the CD8 transmembrane domain^60^ to target NRN1 to the plasma membrane without a GPI anchor. NRN1^116G>E^ migrated differently from wild-type NRN1 on immunoblots (**Fig. 2H**), consistent with altered processing^61, 62^. Both NRN1^116G>E^ and NRN1-ΔGPI were depleted from the membrane fraction and enriched in the conditioned medium relative to wild-type NRN1 (**Fig. 2H**). In contrast, NRN1-TM localized predominantly to the plasma membrane and remained membrane-associated after PI-PLC treatment (**Fig. 2I**). Immunostaining of permeabilized and non-permeabilized cells confirmed these differences in cell-surface localization **(Fig. 2J)**. Functionally, neither NRN1^116G>E^ nor NRN1-ΔGPI enhanced insulin-stimulated AKT phosphorylation (**Figs. 2K,L**). None of the three variants enhanced insulin-stimulated glucose uptake (**Fig. 2L**). Inhibiting GPI-anchor biosynthesis with mannosamine^63^ likewise reduced membrane-associated NRN1 and attenuated the insulin-sensitizing effect of wild-type NRN1 without altering *Nrn1* mRNA (**Supplementary Figs. 3A–C**), phenocopying the anchoring-defective variants (**Supplementary Figs. 3A–C**). Together, these findings indicate that native GPI-anchor processing, and not membrane localization alone, is required for the insulin-sensitizing activity of adipocyte-expressed NRN1.

### Adipose-targeted NRN1 improves adipose function and systemic insulin sensitivity without altering adiposity

Given the marked suppression of NRN1 in visceral adipocytes during obesity, we asked whether restoring it in obese eWAT is sufficient to improve systemic insulin sensitivity. HFD-fed C57BL/6J mice received bilateral intra-eWAT injections of AAV8 encoding NRN1 or control (**Fig. 3A**). This produced robust, depot-selective NRN1 overexpression in eWAT (NRN1^eWAT-OE^) and increased circulating NRN1 without detectable induction of *Nrn1* in other tissues examined (**Figs. 3B–D**).

**Figure 3.**
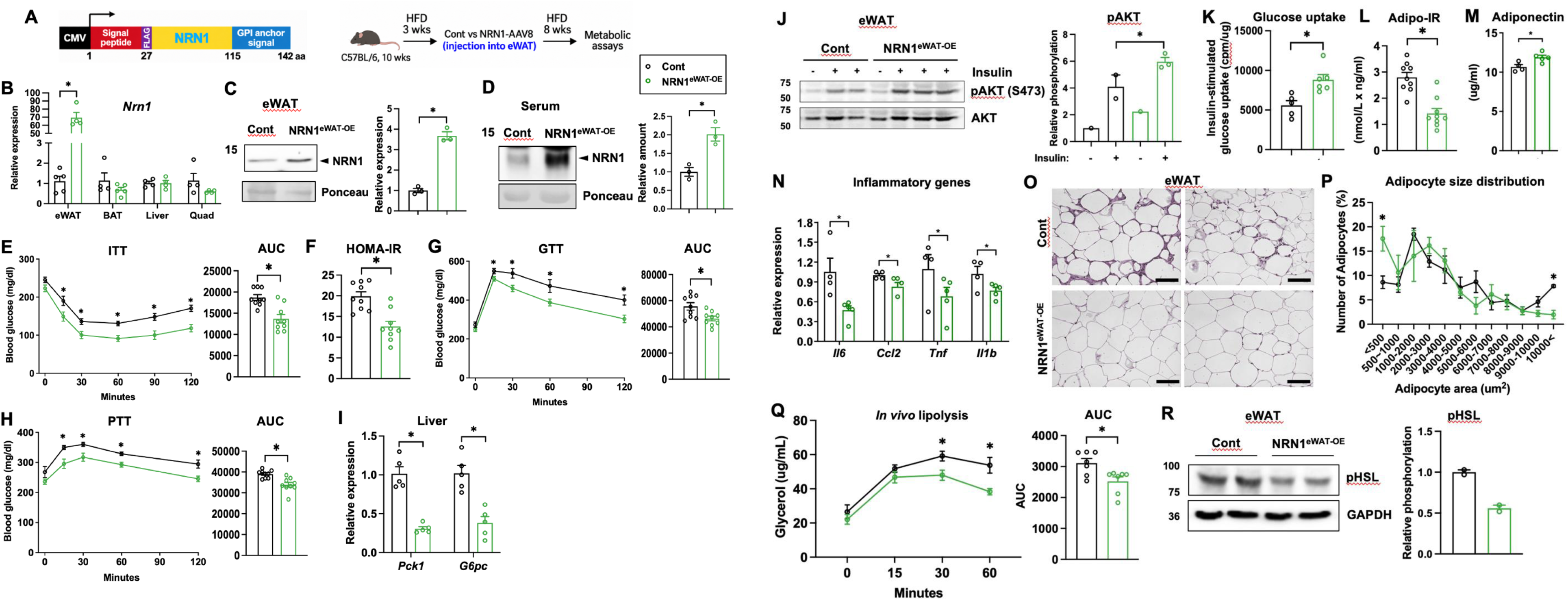
NRN1 overexpression in eWAT improves systemic insulin sensitivity and adipose tissue function in obese mice. **(A)** Schematic of the AAV8-NRN1 construct and experimental timeline. Ten-week-old C57BL/6 mice were fed an HFD for 3 weeks, injected with AAV8-Control (Cont) or AAV8-NRN1 (NRN1^eWAT-OE^) into eWAT, and maintained on the HFD for another 8 weeks before metabolic analyses**. (B)** *Nrn1* mRNA levels in eWAT (*n* = 5 per group), brown adipose tissue (BAT; Cont, *n* = 4; NRN1^eWAT-OE^, *n* = 5), liver and quadriceps muscle (Quad; *n* = 4 per group). **(C)** NRN1 protein levels in eWAT from Cont and NRN1^eWAT-OE^ mice (*n* = 3 per group). **(D)** Circulating NRN1 levels (*n* = 3 per group**). (E)** Insulin tolerance test (ITT) and corresponding area under the curve (AUC) (*n* = 9 per group). **(F)** Homeostatic model assessment of insulin resistance (HOMA-IR) (*n* = 9 per group). **(G)** Glucose tolerance test (GTT) and corresponding AUC in Cont and NRN1^eWAT-OE^ mice (*n* = 9 per group). **(H)** Pyruvate tolerance test (PTT) and corresponding AUC (*n* = 9 per group). **(I)** Hepatic mRNA levels of *Pck1* (*Pepck*) and *G6pc* (*n* = 5 per group). **(J)** AKT phosphorylation in eWAT under basal or insulin-stimulated conditions (basal, *n* = 1 per group; insulin-stimulated Cont, *n* = 2; insulin-stimulated NRN1^eWAT-OE^, *n* = 3). **(K)** Insulin-stimulated glucose uptake in mature adipocytes isolated (Cont, *n* = 5; NRN1^eWAT-OE^, *n* = 6). **(L)** Adipose tissue insulin resistance index (Adipo-IR) (*n* = 9 per group). **(M)** Serum adiponectin levels (Cont, *n* = 4; NRN1^eWAT-OE^, *n* = 5). **(N)** RT–qPCR analysis of inflammatory gene expression in eWAT (Cont, *n* = 4; NRN1^eWAT-OE^, *n* = 5). **(O)** Representative haematoxylin and eosin (H&E)-stained sections of eWAT. Scale bars, 100 µm. **(P)** Quantification of adipocyte number across size bins (*n* = 4 per group). **(Q)** *In vivo* lipolysis assessed by serum glycerol levels and corresponding AUC. **(R)** HSL phosphorylation in eWAT (*n* = 2 per group). Data are presented as mean ± s.e.m. Pairwise comparisons were performed using two-tailed unpaired Student’s *t*-tests.

NRN1^eWAT-OE^ mice showed a greater glucose-lowering response to insulin, improved glucose tolerance and lower HOMA-IR than controls (**Figs. 3E–3G**). Pyruvate tolerance was also improved, accompanied by reduced hepatic expression of the gluconeogenic genes *Pck1* and *G6pc* (**Figs. 3H,I**). These improvements occurred without detectable differences in body weight, body composition or adipose tissue mass (**Supplementary Fig. 4A–C**). Energy expenditure and respiratory exchange ratio (RER), measured by indirect calorimetry, were similar between groups, as were food intake and physical activity (**Supplementary Fig. 4D–L**).

These systemic benefits were accompanied by improved adipose function. eWAT from NRN1^eWAT-OE^ mice showed increased insulin-stimulated AKT phosphorylation and glucose uptake (**Figs. 3J,K**). The adipose insulin resistance index (Adipo-IR), calculated from circulating insulin and free fatty acid concentrations, was lower (**Fig. 3L**), and circulating adiponectin was higher (**Fig. 3M**). eWAT from NRN1^eWAT-OE^ mice also showed lower expression of inflammatory genes and less histological evidence of inflammation (**Fig. 3N,O**). The adipocyte size distribution shifted toward smaller cells, with a lower proportion of hypertrophic adipocytes (**Fig. 3P**). NRN1^eWAT-OE^ also attenuated isoproterenol-stimulated lipolysis and phosphorylation of hormone-sensitive lipase (HSL) in eWAT (**Fig. 3Q,R**). Together, these findings show that increasing NRN1 expression in eWAT is sufficient to enhance local insulin action and improve systemic insulin sensitivity in obese mice without a detectable change in adiposity.

### Systemic NRN1 enhances adipose and whole-body insulin responsiveness

Because circulating NRN1 is reduced in obesity, we asked whether raising systemic NRN1 produces metabolic benefits in obese mice. We fused NRN1 to an Fc domain to increase protein stability and prolong systemic exposure. Intravenous delivery of an AAV8 vector encoding the fusion resulted in predominantly hepatic transduction and increased circulating NRN1–Fc (**Figs. 4A–C**).

**Figure 4.**
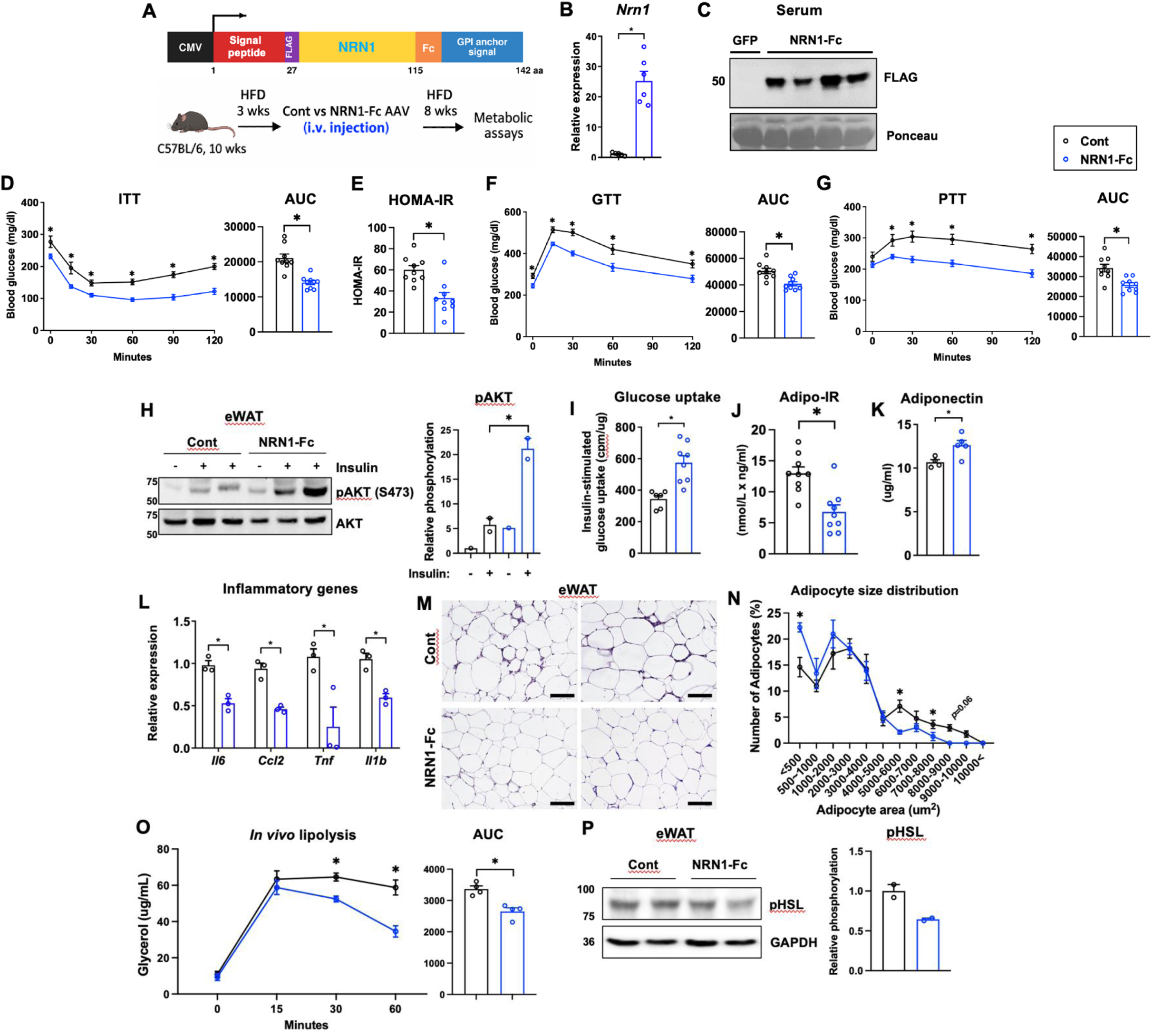
Systemic NRN1–Fc improves insulin sensitivity and adipose tissue function in obese mice. **(A)** Schematic of systemic delivery of AAV8-GFP (Cont) or AAV8-NRN1-Fc (NRN1–Fc) and the experimental timeline. **(B)** *Nrn1* mRNA levels in eWAT (*n* = 6 per group). **(C)** Circulating NRN1 levels (Cont, *n* = 1; NRN1-Fc, *n* = 4). **(D)** ITT and corresponding AUC (*n* = 9 per group). **(E)** HOMA-IR (*n* = 9 per group). **(F)** GTT and corresponding AUC (*n* = 9 per group). **(G)** PTT and corresponding AUC (*n* = 9 per group). **(H)** AKT phosphorylation in eWAT under basal or insulin-stimulated conditions (*n* = 1 and 2 per group, respectively). **(I)** Insulin-stimulated glucose uptake in mature adipocytes (Cont, *n* = 6; NRN1–Fc, *n* = 8). **(J)** Adipo-IR (*n* = 9 per group). **(K)** Serum adiponectin levels (Cont, *n* = 4; NRN1–Fc, *n* = 5). **(L)** RT–qPCR analysis of inflammatory gene expression in eWAT (*n* = 3 per group). **(M)** Representative H&E-stained sections of eWAT. Scale bars, 100 µm. **(N)** Quantification of adipocyte number across size bins (*n* = 4 per group). **(O)** *In vivo* lipolysis and corresponding AUC. **(P)** HSL phosphorylation in eWAT (*n* = 2 per group). Data are presented as mean ± s.e.m. Pairwise comparisons were performed using two-tailed unpaired Student’s *t*-tests.

NRN1–Fc lowered HOMA-IR, enhanced the glucose-lowering response to insulin and reduced glycemic excursions after glucose and pyruvate challenges (**Fig. 4D–G**), without a detectable change in body weight or adiposity (**Supplementary Fig. 5A–C**). In eWAT, NRN1–Fc enhanced insulin signaling and insulin-stimulated glucose uptake (**Fig. 4H,I**). Adipo-IR was lower and circulating adiponectin was higher (**Fig. 4J,K**). eWAT also showed less inflammation and a shift toward smaller adipocytes (**Fig. 4L–N**), along with reduced isoproterenol-stimulated lipolysis and HSL phosphorylation (**Fig. 4O,P**). These adipose phenotypes closely mirrored those observed after eWAT-targeted NRN1 overexpression (**Fig. 3J–R**), supporting eWAT as an NRN1-responsive tissue *in vivo*.

To assess insulin action more directly, we performed tracer-based hyperinsulinemic–euglycemic clamps. Mice expressing NRN1–Fc had markedly lower fasting insulin than controls despite similar fasting glucose concentrations, and plasma insulin remained lower during insulin infusion (**Supplementary Fig. 6A,B**). Despite this lower insulin exposure, whole-body glucose disposal was maintained (**Supplementary Fig. 6C**), and the glucose infusion rate tended to be higher (**Supplementary Fig. 6D**). The plot of whole-body glucose disposal against plasma insulin showed a leftward shift in the NRN1–Fc group (**Supplementary Fig. 6E**), consistent with enhanced whole-body insulin responsiveness. In eWAT, glucose uptake (Rg) was likewise preserved despite lower circulating insulin concentrations (**Supplementary Fig. 6F,G**), consistent with enhanced insulin responsiveness in this depot. Suppression of endogenous glucose appearance (EndoRa) was comparable between groups (**Supplementary Fig. 6H**) and greater in NRN1–Fc mice when assessed relative to plasma insulin (**Supplementary Fig. 6I**). This finding is consistent with the improved pyruvate tolerance in both NRN1 overexpression models (**Figs. 3H and 4G**). Taken together, the clamp data are consistent with enhanced whole-body, adipose and hepatic insulin responsiveness.

### Adipocyte NRN1 protects against diet-induced insulin resistance and impairment of glucose homeostasis

We next asked whether endogenous adipocyte-derived NRN1 is required for metabolic homeostasis. To that end, we generated adipocyte-specific *Nrn1* knockout mice (*Nrn1*^Adi-KO^; **Fig. 5A**) by crossing *Nrn1*^f/f^ mice with *Adipoq*-Cre mice. In knockout mice, *Nrn1* mRNA and NRN1 protein levels were selectively reduced by approximately 50–70% in eWAT, inguinal WAT and BAT (**Figs. 5B,C**). Circulating NRN1 levels were also proportionally reduced (**Fig. 5D**), indicating that adipocytes contribute substantially to the circulating pool.

**Figure 5.**
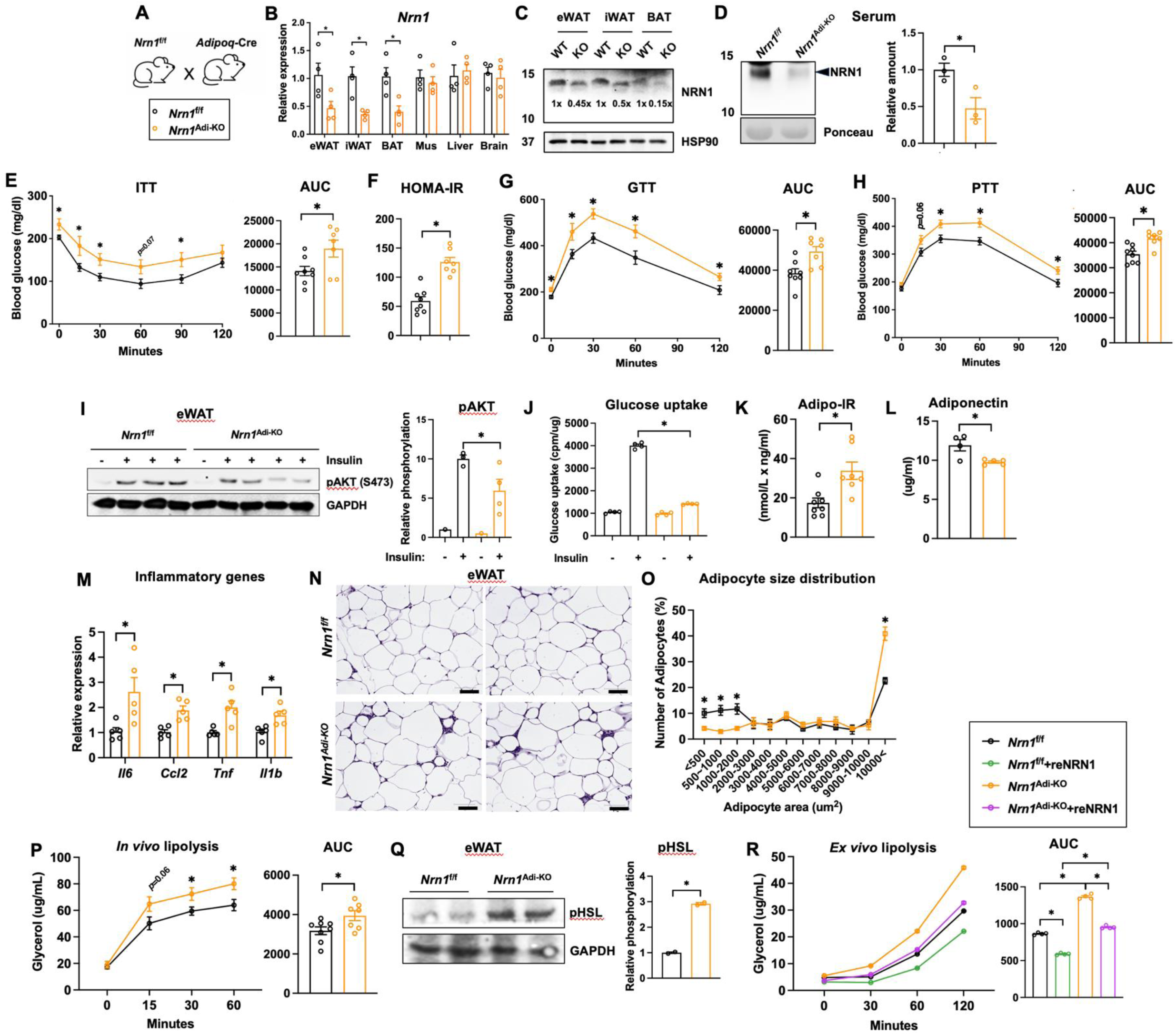
Adipocyte-specific *Nrn1* deletion aggravates insulin resistance and adipose tissue dysfunction in obese mice. **(A,B)** Generation and validation of adipocyte-specific *Nrn1*-knockout (*Nrn1^Adi^*^-KO^) mice. **(A)** Breeding strategy and **(B)** *Nrn1* mRNA levels across tissues (*n* = 4 per group). **(C)** Immunoblot analysis of NRN1 protein in eWAT. **(D)** Circulating NRN1 levels (*n* = 3 per group). **(E)** ITT and corresponding AUC (*Nrn1*^f/f^, *n* = 8; *Nrn1^Adi^*^-KO^, *n* = 7). **(F)** HOMA-IR (*Nrn1*^f/f^, *n* = 8; *Nrn1^Adi^*^-KO^, *n* = 7). **(G)** GTT and corresponding AUC (*Nrn1*^f/f^, *n* = 8; *Nrn1^Adi^*^-KO^, *n* = 7). **(H)** PTT and corresponding AUC (*Nrn1*^f/f^, *n* = 8; *Nrn1^Adi^*^-KO^, *n* = 7). **(I)** AKT phosphorylation in eWAT under basal or insulin-stimulated conditions (basal, *n* = 1 per group; insulin-stimulated *Nrn1*^f/f^, *n* = 3; insulin-stimulated *Nrn1^Adi^*^-KO^, *n* = 4). **(J)** Glucose uptake with fractionated eWAT adipocytes *n* = 4), **(K)** Adipo-IR (*Nrn1*^f/f^, *n* = 8; *Nrn1^Adi^*^-KO^, *n* = 7). **(L)** Serum adiponectin levels (*n* = 5 per group). **(M)** RT–qPCR analysis of inflammatory gene expression in eWAT (*n* = 5 per group). **(N)** Representative H&E-stained sections of eWAT. **(O)** Quantification of adipocyte number across size bins (*n* = 4 per group). **(P)** *In vivo* lipolysis and corresponding AUC (*n* = 7 per group). **(Q)** HSL phosphorylation in eWAT (*n* = 2 per group). **(R)** *Ex vivo* lipolysis in eWAT following overnight treatment with reNRN1 or vehicle, with corresponding AUC (*n* = 4 per group). Data are presented as mean ± s.e.m. Pairwise comparisons were performed using two-tailed unpaired Student’s *t*-tests.

On an HFD, *Nrn1*^Adi-KO^ mice developed greater insulin resistance, poorer glucose tolerance and a larger glycemic response to pyruvate than controls **(Figs. 5E–H**). These systemic glucose homeostatic impairments occurred despite comparable gains in body weight and fat mass. WAT mass, energy expenditure and food intake were also similar between groups (**Supplementary Fig. 7**). Adipocyte *Nrn1* deletion impaired insulin signaling in eWAT and reduced insulin-stimulated glucose uptake in adipocytes from this depot (**Figs. 5I,J**). *Nrn1*^Adi-KO^ mice also had higher Adipo-IR, lower circulating adiponectin and increased eWAT inflammation (**Figs. 5K–N**). The adipocyte size distribution shifted toward larger cells, with a lower proportion of small adipocytes and a higher proportion of hypertrophic adipocytes (**Fig. 5O**), and showed increased lipolysis and HSL phosphorylation were increased (**Figs. 5P,Q**). Notably, *ex vivo* treatment with reNRN1 reduced lipolysis in wild-type eWAT and returned the elevated lipolysis rate in knockout eWAT to control levels (**Fig. 5R**), supporting a direct role for NRN1 in restraining adipose lipolysis. These effects on adipose were opposite to those of eWAT-targeted and systemic NRN1 elevation. Together, these findings from gain- and loss-of-function models identify endogenous adipocyte-derived NRN1 as a protective signal that preserves adipose function and systemic insulin sensitivity during chronic overnutrition, without detectable effect on adiposity.

### BAT thermogenesis does not appear to be a major contributor to NRN1-mediated metabolic protection

BAT-directed NRN1 overexpression was recently reported to enhance thermogenesis^64^. We asked whether BAT thermogenesis is changed in our models and whether any such change contributed to their metabolic phenotypes. eWAT-targeted NRN1 overexpression reduced BAT mass and lipid accumulation (**Supplementary Figs. 8A,F)**. Expression of *Pparg*, *Adrb3*, *Dio2* and *Elovl3* increased modestly in BAT, with no increase in *Ucp1* or *Ppargc1a* expression (**Supplementary Figs. 8D**). Systemic NRN1–Fc preserved BAT morphology without altering BAT mass (**Supplementary Figs. 8B,F**). Adipocyte-specific *Nrn1* deletion modestly increased BAT lipid accumulation without affecting BAT mass or thermogenic gene expression (**Supplementary Figs. 8C,E,G**).

To test whether the metabolic benefits of NRN1 persist when thermogenic demand is minimal, we housed NRN1^eWAT-OE^ and control mice at thermoneutrality for 3 weeks. NRN1^eWAT-OE^ mice remained more insulin sensitive and glucose tolerant than controls, and the differences between groups were similar in magnitude to those observed at room temperature (**Supplementary Fig. 9**). These findings indicate that the metabolic benefits of eWAT-targeted NRN1 overexpression persist when the demand for adaptive thermogenesis is minimized.

In brown adipocyte cultures, the effects of altering NRN1 expression depended on the stage of differentiation. NRN1 overexpression prior to differentiation increased HSL phosphorylation without significantly affecting adipogenesis or UCP1 abundance, whereas *Nrn1* knockdown at this stage impaired brown adipocyte differentiation and reduced UCP1 expression (**Supplementary Figs. 10A,B**). In mature brown adipocytes, NRN1 overexpression, *Nrn1* knockdown, and reNRN1 treatment had little effect on the thermogenic markers examined (**Supplementary Figs. 10C–E**). Together, these findings argue against increased BAT thermogenesis as the principal mediator of NRN1-induced improvements in insulin sensitivity in our animal models.

### Adipocyte FGFR1 is required for the metabolic actions of NRN1

GPI-APs can signal through transmembrane partners, including receptor tyrosine kinases^38-40^. To identify a receptor pathway that might mediate NRN1’s insulin-sensitizing effect, we used a multiplex phospho-kinase array to profile acute signaling in wild-type eWAT explants. Treatment with reNRN1 for 10 min selectively increased phosphorylation of ERK1/2, RSK1/2 and AKT (**Fig. 6A and Supplementary Figs. 11A,B**), a pattern consistent with FGFR pathway activation^65^. Immunoblotting confirmed NRN1-induced ERK1/2 and AKT phosphorylation (**Supplementary Fig. 11C**). Both responses were pronounced in WAT depots and minimal or undetectable in liver and skeletal muscle (**Supplementary Fig. 11D**), identifying WAT as a prominent NRN1-responsive tissue.

**Figure 6.**
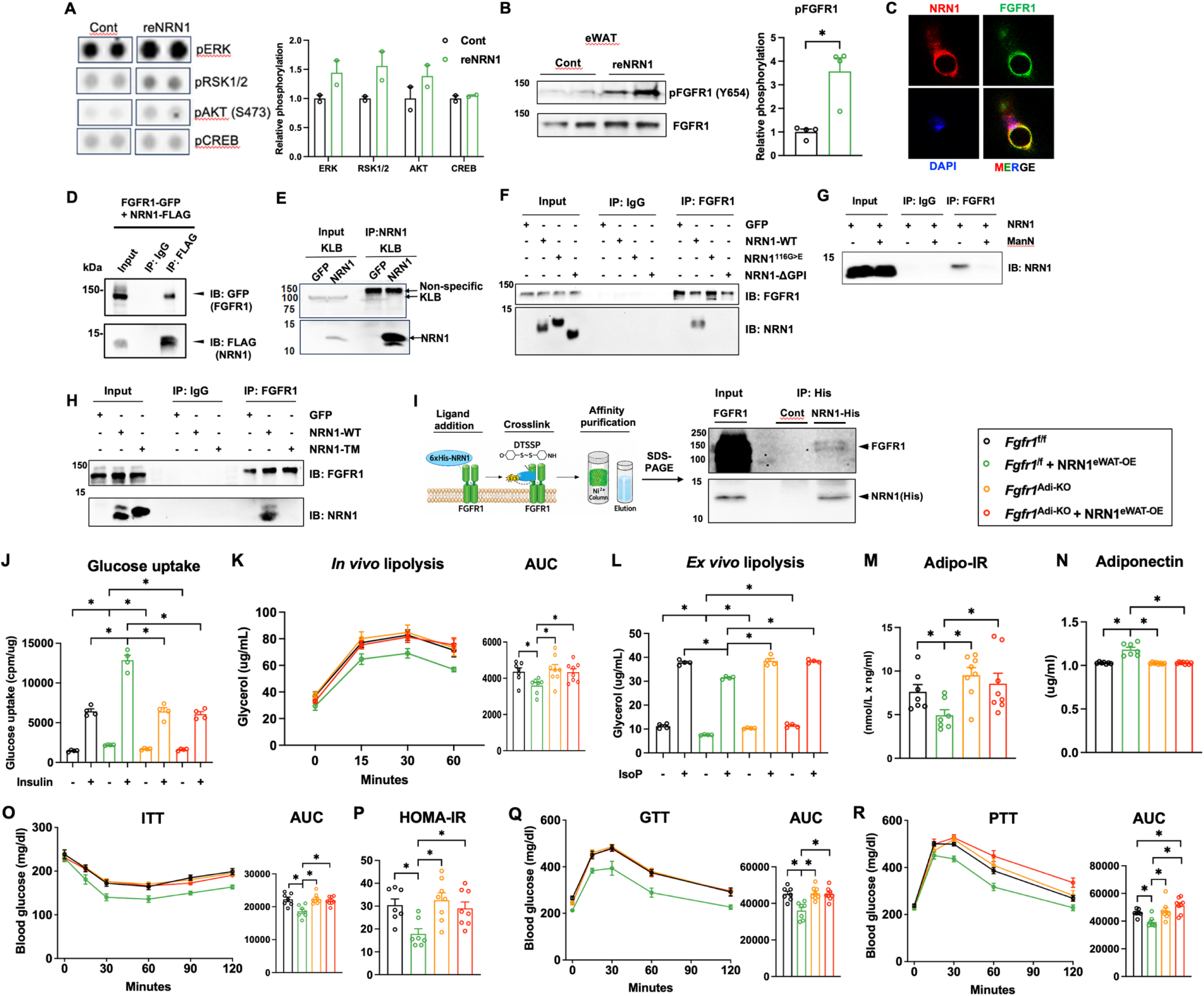
NRN1 engages adipocyte FGFR1 to promote insulin sensitivity. **(A)** Representative duplicate array spots and quantification of selected phosphoproteins— pERK1/2, pRSK1/2, pAKT (Ser473) and pCREB—in eWAT explants treated with Cont or reNRN1 (*n* = 2). **(B)** Representative immunoblots and quantification of total FGFR1 and pFGFR1 (Y654) in wild-type eWAT treated with reNRN1 for 10 min. (*n* = 3). (**C**) Immunofluorescence analysis of NRN1 and FGFR1 in 3T3-L1 adipocytes. Co-IP analyses of the associations of NRN1 with FGFR1 (**D**) and KLB (**E**) in HEK293T cells. **(F)** Co-IP analysis of the association between FGFR1 and NRN1-WT or NRN1^G116<E^, NRN1-ΔGPI in HEK293T cells. **(G)** Co-IP analysis of the NRN1–FGFR1 association in 3T3-L1 adipocytes with or without mannosamine (ManN). (**H)** Co-IP analysis of the association between FGFR1 and NRN1-WT or NRN1-TM in HEK293T cells. **(I)** Schematic and immunoblot analysis of DTSSP-mediated NRN1–FGFR1 crosslinking and co-IP in 3T3-L1 adipocytes. *Ex vivo* insulin stimulated glucose uptake with eWAT (**J**) (*n* = 4 per group), *in vivo* lipolysis assay with corresponding AUC (**K**) (*n* = 7 per group), *ex vivo* lipolysis assay in eWAT, with corresponding AUC (**L**) (*n* = 4 per group), Adipo-IR (**M**) (*n* = 7 per group), and serum adiponectin levels (**N**), ITT with corresponding AUC (**O**), HOMA-IR (**P**), GTT with corresponding AUC (**Q**) (*n* = 7 per group), PTT with corresponding AUC (**R**) (*n* = 7 per group), in *Fgfr1^f/f^*, NRN1^eWAT-OE^, *Fgfr1*^Adi-KO^ and *Fgfr1*^Adi-KO^ + NRN1^eWAT-OE^ mice. Data are presented as mean ± s.e.m. Pairwise comparisons were performed using two-tailed unpaired Student’s *t*-tests. Two-way ANOVA was used for datasets with two independent variables.

A prior neuronal study implicated NRN1–FGFR signaling in axonal branching^66^. FGF1 and FGF21 both enhances systemic insulin sensitivity in obese mice^41, 67^, and signal through FGFR1 in adipocytes^43, 68^. We therefore focused on FGFR1-insulin sensitizing effect of NRN1. reNRN1 increased FGFR1 tyrosine phosphorylation in eWAT explants within 10 min (**Fig. 6B**), with a stronger response in eWAT than in the other tissues examined (**Supplementary Fig. 11D**). NRN1 and FGFR1 colocalized at the adipocyte surface (**Fig. 6C**) and co-immunoprecipitated (**Fig. 6D).** The same assay detected no association between NRN1 and β-Klotho (KLB) (**Fig. 6E**). Notably, FGFR1 association was undetectable with either NRN1^116>E^ or NRN1-ΔGPI and with cell-expressed wild-type NRN1 after inhibition of GPI-anchor biosynthesis (**Fig. 6F,G**). NRN1-TM also failed to co-precipitate with FGFR1 despite its plasma-membrane localization (**Fig. 6H**). Thus, native GPI-anchor processing is necessary for cell-expressed NRN1 to associate with FGFR1. Crosslinking with the membrane-impermeable reagent 3,3′-dithiobis(sulfosuccinimidyl propionate) (DTSSP) further supported close proximity between NRN1 and FGFR1 at the adipocyte surface (**Fig. 6I**). Collectively, these data support engagement of adipocyte FGFR1 by NRN1 and NRN1-dependent FGFR1 activation.

We next tested whether FGFR1 mediates the insulin-sensitizing effect of NRN1 in cultured adipocytes. Pharmacological FGFR inhibition abolished the ability of NRN1 to enhance insulin-stimulated glucose uptake in adipocytes cultured with or without TNF (**Supplementary Fig. 12A**). CRISPR–Cas9-mediated *Fgfr1* knockdown largely eliminated this response (**Supplementary Fig. 12B**). By contrast, *Fgfr2* knockdown preserved the response to NRN1 under control conditions and modestly enhanced it under TNF challenge (**Supplementary Fig. 12C**), supporting a specific requirement for FGFR1.

To test this requirement *in vivo*, we overexpressed NRN1 in eWAT of adipocyte-specific *Fgfr1*-knockout mice (*Fgfr1*^AKO^) and floxed littermate controls (**Supplementary Figs. 13A,B**). Among HFD-fed mice receiving the control vector, floxed controls and *Fgfr1*^AKO^ mice showed comparable HOMA-IR and responses to insulin, glucose and pyruvate challenges (**Fig. 6N–Q**), consistent with previous reports^68, 69^. In floxed mice, eWAT-targeted NRN1 overexpression improved insulin, glucose and pyruvate tolerance and lowered HOMA-IR relative to the control vector. Strikingly, these systemic benefits were largely absent in *Fgfr1*^AKO^ mice (**Fig. 6N–Q**). Body weight and body composition were similar across all groups (**Supplementary Fig. 13C**). Together with the biochemical, pharmacological and cellular evidence, these epistasis experiments indicate that adipocyte FGFR1 is required for the local adipose and systemic insulin-sensitizing effects of NRN1.

### NRN1–FGFR1 signaling promotes insulin sensitivity through a PPARγ-dependent program

To identify pathways that mediate the insulin-sensitizing effects of NRN1–FGFR1 signaling, we integrated eWAT transcriptomes from NRN1^eWAT-OE^ and *Nrn1*^Adi-KO^ mice. We identified 439 genes positively regulated by NRN1 (upregulated by gain and downregulated by loss of function; **Fig. 7A**) and 425 genes regulated in the opposite direction (**Supplementary Fig. 14**). PPARγ signaling was among the most strongly enriched pathways in the positively regulated gene set (**Fig. 7A and Supplementary Fig. 14**). Given PPARγ’s central role in adipose metabolism and its reported links to FGF/FGFR signaling^45, 49-51^, we focused on PPARγ as a candidate downstream mediator.

**Figure 7.**
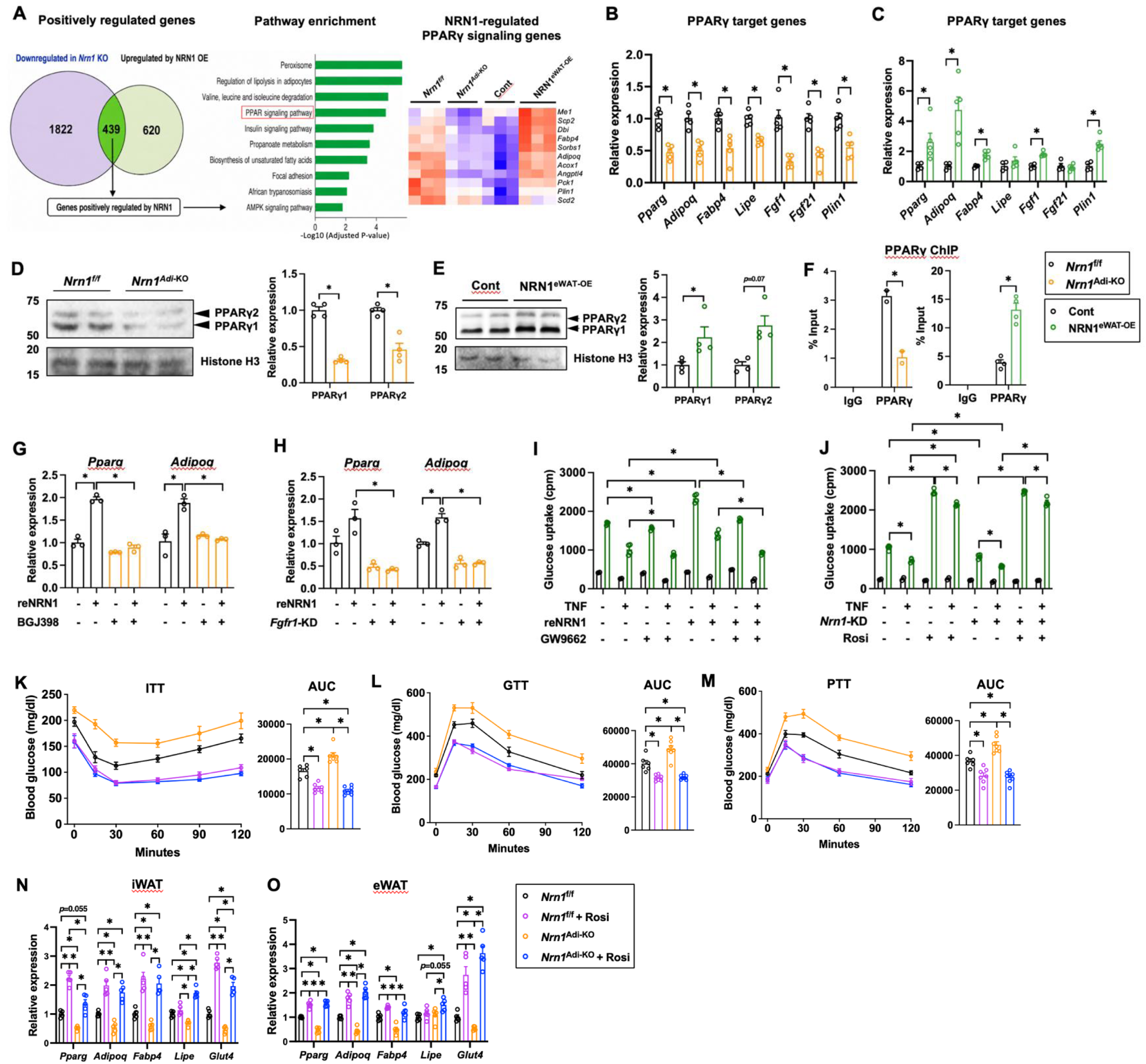
Adipocyte NRN1–FGFR1 signaling promotes insulin sensitivity through PPARγ. **(A)** Identification of NRN1-positively regulated genes based on their downregulation in *Nrn1^Adi^*^-KO^ eWAT and upregulation in NRN1^eWAT-OE^ eWAT, followed by KEGG pathway analysis and heatmap visualization of NRN1-regulated PPAR signaling genes. **(B, C)** mRNA levels of PPARγ target genes in eWAT from *Nrn1^Adi^*^-KO^ **(B)** and NRN1^eWAT-OE^ (**C**) mice and their respective controls (*n* = 5 per group). **(D, E)** Immunoblot analysis of PPARγ in eWAT from *Nrn1^Adi^*^-KO^ **(D)** and NRN1^eWAT-OE^ **(E)** mice and their respective controls (*n* = 4 per group). **(F)** ChIP–qPCR analysis of PPARγ occupancy in eWAT from *Nrn1^Adi^*^-KO^ and NRN1^eWAT-OE^ mice and their respective controls (*n* = 2 and *n* = 4 from *Nrn1^Adi^*^-KO^ and NRN1^eWAT-OE^ groups, respectively). *Pparg* and *Adipoq* mRNA levels in 3T3-L1 adipocytes treated with BGJ398 (**G**) or transduced with control or *Fgfr1*-knockdown lentivirus **(H)**, followed by reNRN1 treatment (*n* = 3). Glucose uptake in 3T3-L1 adipocytes treated with or without TNFα following PPARγ antagonism in the presence or absence of reNRN1 **(I)** or PPARγ agonism in control and *Nrn1*-knockdown cells **(J)**. (*n* = 2 for basal conditions and *n* = 4 for insulin-stimulated conditions). ITT **(K)**, GTT **(L)** and PTT **(M)**, with corresponding, with corresponding AUCs, in *Nrn1*^f/f^ and *Nrn1^Adi^*^-KO^ mice treated with vehicle or rosiglitazone (Rosi) (*Nrn1*^f/f^, *n* = 6; *Nrn1*^f/f^ + Rosi and *Nrn1^Adi^*^-KO^, *n* = 7; *Nrn1^Adi^*^-KO^ + Rosi, *n* = 8). mRNA levels of PPARγ target genes in inguinal WAT (iWAT; **N**) and eWAT **(O)** in the same four groups (*n* = 5 per group). Data are presented as mean ± s.e.m. Pairwise comparisons were performed using two-tailed unpaired Student’s *t*-tests. Two-way ANOVA was used for datasets with two independent variables.

Independent qPCR analyses confirmed higher expression of *Pparg*, *Adipoq* and selected PPARγ-responsive metabolic genes with NRN1 overexpression and lower expression with *Nrn1* deficiency (**Fig. 7B,C**). *Fgf1* expression followed the same pattern, and *Nrn1* deficiency also reduced *Fgf21* expression (**Fig. 7B,C**). Because *Fgf1* and *Fgf21* are PPARγ targets that encode ligands that signal through FGFR1^45, 49^, these changes raise the possibility that PPARγ-dependent FGF expression reinforces FGFR1 signaling. PPARγ protein abundance also increased with NRN1 overexpression and decreased with *Nrn1* deficiency (**Fig. 7D,E**). PPARγ occupancy at the *Pparg* locus followed the same pattern (**Fig. 7F**), consistent with positive autoregulation of *Pparg*. Pharmacological FGFR inhibition and *Fgfr1* knockdown each blocked NRN1-induced expression of *Pparg* and *Adipoq* in adipocytes (**Fig. 7G,H**), placing FGFR1 upstream of this transcriptional response.

We next tested whether PPARγ activity is required for NRN1 to enhance insulin responsiveness. The PPARγ antagonist GW9662 abolished the increase in insulin-stimulated glucose uptake elicited by NRN1 in adipocytes (**Fig. 7I**). The PPARγ agonist rosiglitazone (Rosi) restored insulin-stimulated glucose uptake in *Nrn1*-deficient adipocytes (**Fig. 7J**). This rescue extended to the whole-animal: Rosi also largely corrected the metabolic defects *in vivo*. After 3 weeks of treatment, *Nrn1*^Adi-KO^ mice and Rosi-treated controls showed comparable glycemic responses in the insulin, glucose and pyruvate tolerance tests (**Figs. 7K,M**). Rosi also substantially restored expression of NRN1-regulated PPARγ-responsive genes in knockout WATs (**Figs. 7N,O**). Together, these findings place PPARγ downstream of NRN1–FGFR1 signaling in adipocytes and show that pharmacological PPARγ activation can largely compensate for adipocyte *Nrn1* loss *in vivo*.

## Discussion

A central question in obesity biology is which adipocyte signals protect insulin action and are diminished as adipose tissue expands. Here, we identify NRN1 as an obesity-suppressed, adipocyte-derived signal that helps preserve white adipose tissue function and systemic insulin sensitivity. Our findings link obesity-associated DNMT3A to reduced NRN1 and identify adipocyte FGFR1–PPARγ signaling as a mechanism through which NRN1 sustains insulin responsiveness.

We previously showed that adipose DNMT3A is elevated in obesity and contributes causally to insulin resistance^8^. DNMT3A occupied the *Nrn1* promoter, and increasing or reducing DNMT3A had opposing effects on promoter methylation and *Nrn1* expression. These findings support *Nrn1* as a likely direct target of DNMT3A. In obese mice, increased *Nrn1* promoter methylation in eWAT coincided with lower NRN1 expression, less NRN1 release from eWAT explants and lower circulating NRN1 levels. Adipocyte-specific *Nrn1* deletion lowered circulating NRN1 levels, indicating adipocytes as a source of the circulating pool. Similarly, in humans, adipocyte NRN1 expression and plasma NRN1 concentrations were each inversely associated with BMI. Together, these observations are consistent with a model in which DNMT3A-mediated repression contributes to the obesity-associated decline in adipocyte-derived NRN1, thereby reducing a protective signal that helps sustain insulin responsiveness.

Gain- and loss-of-function studies identify WAT as a major site of NRN1 action. Both eWAT-targeted NRN1 overexpression and circulating NRN1–Fc enhanced adipose insulin responsiveness and raised circulating adiponectin, while reducing inflammation, adipocyte hypertrophy and stimulated lipolysis. Adipocyte-specific *Nrn1* deletion produced reciprocal phenotypes without altering energy expenditure or fat mass. Because NRN1–Fc lowered circulating insulin, the clamp studies cannot be compared between groups at matched insulin exposure. Nevertheless, preserved whole-body glucose disposal and eWAT glucose uptake at lower insulin concentrations indicate enhanced insulin responsiveness. This interpretation is further strengthened by reciprocal effects of NRN1 gain and loss on insulin-stimulated glucose uptake in eWAT explants under controlled insulin exposure, as well as by direct insulin sensitization by NRN1 in isolated mouse and human adipocytes.

Our findings support NRN1 as a non-FGF activator of adipocyte FGFR1 signaling. NCAM provides a precedent for FGFR engagement by a non-FGF protein in the nervous system^40^. NRN1 colocalized with FGFR1 at the adipocyte surface and co-immunoprecipitated with the receptor. Cell-surface crosslinking further supported their proximity. Acute reNRN1 treatment increased FGFR1 tyrosine phosphorylation and activated ERK–RSK and AKT signaling, with the strongest responses in eWAT explants. Pharmacological FGFR inhibition abolished NRN1-mediated insulin sensitization in cultured adipocytes, and *Fgfr1* knockdown largely eliminated this response. In mice, adipocyte-specific *Fgfr1* deletion largely abolished the adipose and systemic benefits of eWAT-targeted NRN1 overexpression.

For adipocyte-expressed NRN1, detectable NRN1–FGFR1 association and insulin-sensitizing activity depended on native GPI-anchor processing. Replacing the GPI-anchoring signal with the CD8 transmembrane domain preserved cell-surface localization, yet NRN1-TM failed to associate detectably with FGFR1 or enhance insulin-stimulated glucose uptake. Soluble reNRN1 enhanced glucose uptake in recipient adipocytes, as did conditioned medium from NRN1-overexpressing HEK293T cells, which lack endogenous NRN1. The GPI-disrupting variants were enriched in conditioned medium, yet expression of either variant failed to enhance insulin responsiveness. Differences in processing, molecular form or effective concentration could explain this result. The relative contributions of soluble and membrane-associated NRN1 *in vivo* remain unresolved.

PPARγ emerged as a key effector linking NRN1–FGFR1 signaling to adipocyte function. FGFR inhibition and *Fgfr1* knockdown each blocked NRN1-dependent induction of *Pparg* and *Adipoq*. PPARγ abundance, occupancy at the *Pparg* locus and expression of PPARγ-responsive genes increased with NRN1 overexpression and decreased with *Nrn1* loss. PPARγ antagonism blocked NRN1-mediated insulin sensitization in adipocytes, and Rosi restored insulin-stimulated glucose uptake in *Nrn1*-deficient adipocytes and largely corrected the metabolic impairments in knockout mice. The reciprocal changes in circulating adiponectin and eWAT inflammation are consistent with PPARγ’s established roles in promoting adiponectin production and restraining adipose inflammation^70, 71^. Whether PPARγ also mediates NRN1-dependent suppression of lipolysis remains open, given its context-dependent effects on this process^72-75^.

The hepatic effects of NRN1 may partly reflect improved adipose function and reduced lipolysis. NRN1–Fc-expressing mice maintained suppression of endogenous glucose appearance at lower plasma insulin concentrations than controls, consistent with enhanced hepatic insulin responsiveness. NRN1 elicited little canonical signaling in the liver and did not detectably activate insulin signaling in hepatocytes, consistent with low hepatic FGFR1 expression. In eWAT explants, reNRN1 reduced lipolysis in wild-type tissue and corrected the elevated lipolysis associated with *Nrn1* deficiency. By restraining adipose lipolysis, NRN1 could reduce the delivery of glycerol and fatty acids to the liver. This would limit glycerol availability for gluconeogenesis and could reduce hepatic acetyl-CoA production from fatty acids^76, 77^, providing a plausible link between adipose NRN1 signaling and hepatic glucose production.

BAT-directed NRN1 overexpression has been reported to enhance thermogenesis^64^. In our models, changes in BAT morphology and gene expression were modest and varied across interventions, and NRN1 had little effect on the thermogenic program of mature brown adipocytes. The improvements in insulin sensitivity and glucose tolerance after eWAT-targeted NRN1 overexpression persisted at thermoneutrality. Moreover, neither eWAT-targeted overexpression nor adipocyte *Nrn1* deletion altered whole-body energy expenditure. These findings argue against adaptive thermogenesis as the principal mediator of the metabolic benefits observed here.

The adiposity-independent insulin-sensitizing effects of NRN1 raise therapeutic interest. Direct PPARγ agonists such as rosiglitazone improve insulin sensitivity but can cause weight gain and fluid retention and increase the risk of heart failure^10, 78, 79^. Pharmacological FGF1 improves glucose homeostasis through adipose FGFR1 in obese mice^41, 76^, and engineered FGF21 analogues provide a precedent for therapeutically targeting endocrine FGF signaling^80-82^. In our models, NRN1 engaged a PPARγ-dependent program through adipocyte FGFR1 and improved insulin sensitivity despite persistent obesity. Whether NRN1-based approaches can sustain this benefit in humans with adequate tissue selectivity and without the fluid retention and cardiac risks associated with direct PPARγ agonism remains to be established.

Together, our findings identify NRN1 as an obesity-suppressed adipokine that undergoes GPI anchoring during maturation and define a DNMT3A–NRN1–FGFR1–PPARγ pathway that supports adipocyte function and systemic insulin sensitivity. Restoring NRN1 signaling may improve insulin responsiveness without altering adiposity, a possibility that warrants further testing.

---

**Supplementary Figure 1. NRN1 expression is reduced in *ob/ob* mice and increases during mouse and human adipocyte differentiation. (A)** Relative *Nrn1* mRNA levels in eWAT from *ob/+* control and *ob/ob* mice (*n* = 3 per group). **(B)** Relative *Nrn1* mRNA levels in confluent preadipocytes (PreAds) and mature 3T3-L1 adipocytes (Ads) (*n* = 3). **(C)** Relative *NRN1* mRNA levels in confluent preadipocytes (PreAds) and mature human SGBS adipocytes (Ads) (*n* = 3). Data are presented as mean ± SEM. Statistical significance was determined by an unpaired two-tailed Student’s *t*-test. \**P* < 0.05.

**Supplementary Figure 2. Validation of NRN1 overexpression and knockdown in mouse and human adipocytes. (A)** Representative immunoblots and quantification of NRN1 protein in 3T3-L1 adipocytes transduced with GFP-control or NRN1-overexpression lentivirus, or with control or *Nrn1*-knockdown lentivirus. HSP90 served as the loading control (*n* = 3). **(B)** Relative *NRN1* mRNA levels in human SGBS adipocytes transduced with control or *NRN1*-knockdown lentivirus (*n* = 3). Data are presented as mean ± s.e.m. Pairwise comparisons were performed using two-tailed unpaired Student’s *t*-tests. \**P* < 0.05.

**Supplementary Figure 3. Inhibition of GPI-anchor biosynthesis attenuates the insulin-sensitizing effect of NRN1 in adipocytes. (A)** Immunoblot analysis of membrane-associated and released NRN1 in 3T3-L1 adipocytes treated with or without the GPI-anchor biosynthesis inhibitor D-mannosamine (ManN; *n* = 2). **(B)** Relative *Nrn1* mRNA levels in 3T3-L1 adipocytes treated with or without ManN (*n* = 2). **(C)** Glucose uptake in 3T3-L1 adipocytes transduced with control or NRN1-overexpression lentivirus and treated with or without TNF and ManN under basal or insulin-stimulated conditions (*n* = 2 and 4, respectively). Data are presented as mean ± s.e.m. Pairwise comparisons were performed using two-tailed unpaired Student’s *t*-tests.

**Supplementary Figure 4. NRN1eWAT^-OE^ mice show no discernible changes in energy homeostasis. (A)** Body-weight trajectory during HFD feeding; the arrow indicates AAV administration. **(B)** Body composition, expressed as percentages of body weight, and **(C)** tissue weights in control and NRN1^eWAT-OE^ mice after HFD feeding (*n* = 7 per group). **(D)** Relationship between energy expenditure and body weight, analyzed by ANCOVA with body weight as the covariate. **(E)** Average energy expenditure. Oxygen consumption (VO₂) over the light–dark cycle (**F**) and average VO₂ (**G**). Carbon dioxide production (VCO₂) over the light–dark cycle (**H**) and average VCO₂ (**I**). **(J)** Food intake, **(K)** respiratory exchange ratio (RER), and **(L)** locomotor activity. Indirect calorimetry was performed at RT (*n* = 7 per group). Data are presented as mean ± s.e.m. The relationship between energy expenditure and body weight in **D** was analyzed by ANCOVA. Other pairwise comparisons were performed using two-tailed unpaired Student’s *t*-tests.

**Supplementary Figure 5. NRN1–Fc-treated mice show no discernible changes in body weight or adiposity. (A)** Body-weight gain during HFD feeding. **(B)** Body composition normalized to body weight and **(C)** tissue weights after HFD feeding.

**Supplementary Figure 6. NRN1 enhances eWAT insulin responsiveness while preserving whole-body glucose disposal at lower insulin concentrations.** Fasting blood glucose (**A**), plasma insulin under fasting and clamp conditions (**B**), total glucose disappearance rate (Rd; **C**), and glucose infusion rate (GIR; **D**) in control and NRN1–Fc-treated mice. (**E**) Relationship between Rd and plasma insulin; lines connect group means. eWAT glucose uptake rate (Rg) during the clamp (**F**) and its relationship with plasma insulin (**G**); lines represent group-specific linear regressions. Endogenous glucose appearance rate (EndoRa), reflecting hepatic glucose production (**H**), and its relationship with plasma insulin (**I**); lines connect group means. Unless otherwise indicated, Cont, *n* = 9; NRN1–Fc, *n* = 11. Mice lacking paired insulin measurements for Rd, Rg, or EndoRa were excluded from the corresponding analyses in **E, G**, and **I**. Data are presented as mean ± s.e.m. Statistical significance was assessed using two-tailed unpaired Student’s *t*-tests. *P* < 0.05.

**Supplementary Figure 7. Adipocyte-specific *Nrn1* deletion does not alter body composition or energy homeostasis. (A)** Body-weight trajectory during HFD feeding. **(B)** Body composition, expressed as a percentage of body weight. **(C)** iWAT and eWAT white adipose tissue weights. **(D)** Relationship between energy expenditure and body weight, analyzed by ANCOVA with body weight as the covariate. **(E)** Average energy expenditure. VO₂ across the light–dark cycle (**F**) and average VO₂ (**G**). VCO₂ across the light–dark cycle (**H**) and average VCO₂ (**I**). **(J)** Food intake, **(K)** RER, and **(L)** locomotor activity. Indirect calorimetry was performed at 25 °C. For **A** and **C**, *n* = 6 per group; for **B**, *n* = 4 *Nrn1*^f/f^ and 5 *Nrn1^Adi^*^-KO^ mice; and for **D–L**, *n* = 4 per group. Data are presented as mean ± s.e.m. The analysis in **D** was performed by ANCOVA. Other pairwise comparisons were performed using two-tailed unpaired Student’s *t*-tests.

**Supplementary Figure 8. NRN1 modulates BAT mass and lipid accumulation without broadly activating the canonical thermogenic program.** BAT weights in Cont and NRN1^eWAT-OE^ mice **(A)**, Cont and NRN1-Fc-treated mice **(B)**, and *Nrn1*^f/f^ and 5 *Nrn1^Adi^*^-KO^ mice **(C)** (control groups in **A** and **B**, *n* = 3; all other groups, *n* = 5). Relative mRNA levels of *Ucp1*, *Pgc1a*, *Pparg*, *Adrb3*, *Dio2*, and *Elovl3* in BAT from Cont and NRN1^eWAT-OE^ mice **(D)** and from *Nrn1*^f/f^ and 5 *Nrn1^Adi^*^-KO^ mice **(E)** (*n* = 4 per group). Representative H&E-stained BAT sections from control, NRN1^eWAT-OE^, and NRN1-Fc-treated mice **(F)** and from *Nrn1*^f/f^ and 5 *Nrn1^Adi^*^-KO^ mice **(G)**. Scale bars, 100 µm. Data are presented as mean ± s.e.m. Pairwise comparisons were performed using two-tailed unpaired Student’s *t*-tests. Exact *P* values are shown where indicated; \**P* < 0.05.

**Supplementary Figure 9. The metabolic benefits of eWAT-targeted NRN1 overexpression persist at thermoneutrality.** Cont and NRN1^eWAT-OE^ mice (*n* = 7 per group) were evaluated at room temperature (RT; **A,B**) and following 3–4 weeks of thermoneutral (TN) housing **(C–E)**. **(A,C)** ITTs with corresponding AUCs. **(B,D)** GTTs with corresponding AUCs. **(E)** HOMA-IR. Data are presented as mean ± s.e.m. Pairwise comparisons were performed using two-tailed unpaired Student’s *t*-tests. Exact *P* values are shown where indicated; \**P* < 0.05.

**Supplementary Figure 10. Effects of NRN1 gain and loss of function on adrenergic and thermogenic signaling in brown adipocytes.** Experimental design, representative immunoblots, and quantification in immortalized brown preadipocytes transduced 3 days before differentiation with GFP-control or NRN1-overexpression lentivirus **(A)** or control or *Nrn1*-knockdown lentivirus **(B)** and collected on differentiation day 6. **(C,D)** Corresponding analyses in mature brown adipocytes transduced on differentiation day 6 and collected 3 days later. **(E)** Immunoblot analysis and quantification in mature brown adipocytes treated with or without reNRN1 for 24 h and stimulated with IsoP during the final hour where indicated. *n* = 2 per condition. Data are mean ± s.e.m.; two-tailed unpaired Student’s *t*-tests. *P* < 0.05.

**Supplementary Figure 11. NRN1 selectively induces an FGFR-associated signaling signature in eWAT.** Representative phospho-kinase arrays **(A)** and quantification normalized to reference spots **(B)** in vehicle (Cont)- and reNRN1-treated wild-type eWAT explants (*n* = 2). Target identities are listed at right. **(C)** Immunoblot analysis and quantification of AKT and ERK phosphorylation in eWAT explants. **(D)** Immunoblot analysis of FGFR1-related signaling proteins in the indicated metabolic tissues treated with or without reNRN1 for 10 min. Ponceau staining served as a loading control. Data are mean ± s.e.m.; two-tailed unpaired Student’s *t*-tests.

**Supplementary Figure 12. FGFR1 is required for the insulin-sensitizing effects of NRN1 in adipocytes.** Glucose uptake in 3T3-L1 adipocytes treated with vehicle or the FGFR inhibitor BGJ398 (**A**), transduced with control or *Fgfr1*-knockdown lentivirus (**B**), or transduced with control or *Fgfr2*-knockdown lentivirus (**C**). Cells were treated with or without TNF and reNRN1 and assessed under basal or insulin-stimulated conditions (*n* = 2 and 4, respectively).

**Supplementary Figure 13. Validation of adipocyte-specific *Fgfr1* deletion and body composition following eWAT-targeted NRN1 overexpression. Immunoblot analysis and quantification of FGFR1 (A)** and NRN1 **(B)** in tissues from *Fgfr1^f/f^* and *Fgfr1*^Adi-KO^ mice (*n* = 2 per group). **(C)** Body composition measured by EchoMRI in *Fgfr1^f/f^*, *Fgfr1^f/f^* + NRN1^eWAT-OE^, *Fgfr1*^Adi-KO^, and *Fgfr1*^Adi-KO^ + NRN1^eWAT-OE^ mice (*n* = 7 for *Fgfr1^f/f^* groups and *n* = 8 for *Fgfr1*^Adi-KO^ groups). Data are mean ± s.e.m. Two-tailed unpaired Student’s *t*-tests were used for pairwise comparisons, and two-way ANOVA was used for analyses involving genotype and NRN1 overexpression.

**Supplementary Figure 14. Transcriptomic identification of genes negatively regulated by NRN1 in eWAT. (A)** Venn diagram showing the overlap between genes upregulated in eWAT from *Nrn1^Adi^*^-KO^ mice and genes downregulated in eWAT from NRN1^eWAT-OE^ mice, identifying 425 genes negatively regulated by NRN1. **(B)** KEGG pathway enrichment analysis of the 425 overlapping genes. Bars represent −log10-adjusted *P* values.

## Methods

### SVF and mature adipocyte isolation

Adipose tissue pads were excised, minced in digestion buffer containing 2 mg/mL collagenase type II, 10 mM CaCl₂, and 3% BSA in PBS, and incubated at 37°C for 30 min with shaking. The digested tissue was passed through a 100-µm cell strainer and centrifuged at 300 × *g* for 5 min at room temperature. The upper layer containing mature adipocytes was collected, while the pelleted SVF was treated with red blood cell lysis buffer for 5 min at room temperature. The mature adipocyte and SVF fractions were then passed through 40-µm cell strainers, centrifuged at 500 × *g* for 5 min at room temperature, and resuspended in PBS containing 3% BSA for washing. Following an additional centrifugation at 500 × *g* for 5 min at room temperature, the mature adipocyte and SVF fractions were collected for subsequent experiments.

### Cell supernatants and serum concentration

Serum samples and supernatants filtered through a 0.45-µm membrane were diluted with five volumes of RIPA buffer and incubated on ice for 10 min, with vortexing every 5 min. The samples were then centrifuged at 10,000 × *g* for 10 min at 4°C to remove insoluble material. The resulting supernatants were concentrated using centrifugal filters with a 10-kDa molecular-weight cutoff at 5,000 × *g* for 30 min at 4°C. PBS containing protease inhibitors was added to the concentrated samples, followed by an additional centrifugation at 5,000 × *g* for 30 min at 4°C. The retentates remaining in the filters were collected for subsequent experiments.

### Animal models

*Nrn1*^Adi-KO^ and *Fgfr1*^Adi-KO^ were generated by crossing Adipoq-Cre mice (JAX stock no. 028020) with *Nrn1*^f/f^ mice (JAX stock no. 034075) or *Fgfr1^f/f^* mice (JAX stock no. 007671). For systemic NRN1 elevation, AAV8 encoding NRN1–Fc or GFP (4.5x10^10^ vg/mouse) was administered via tail-vein injection. For eWAT-selective NRN1 overexpression, AAV8 encoding *Nrn1* or GFP (3x10^10^ vg/mouse) was injected bilaterally into the eWAT depots. Mice were maintained under a 12-h light/12-h dark cycle with *ad libitum* access to food and water. Diet-induced obesity was established by feeding C57BL/6 mice a HFD diet containing 60% of calories from fat (Research Diets, D12492). Experimental groups were compared with age-matched littermate controls. All animal procedures were performed in accordance with institutional guidelines and were approved by the Institutional Animal Care and Use Committee.

### Metabolic phenotyping

For ITTs, mice were fasted for 6 h and injected intraperitoneally with insulin at 0.5 U kg⁻¹ for chow-fed mice or 0.8–1.2 U kg⁻¹ for HFD-fed mice. For GTTs, mice were fasted for 6 h and injected intraperitoneally with glucose at 1 g kg⁻¹ for chow-fed mice or 1.5 g kg⁻¹ for HFD-fed mice. For PTTs, HFD-fed mice were fasted for 6 h and injected intraperitoneally with pyruvate at 1.2–1.5 g kg⁻¹. Blood glucose concentrations were measured using a Contour NEXT glucometer and compatible test strips at 0, 15, 30, 60, 90, and 120 min for ITTs and at 0, 15, 30, 60, and 120 min for GTTs and PTTs. Metabolic parameters were assessed by open-circuit indirect calorimetry using Oxymax chambers within the Comprehensive Laboratory Animal Monitoring System (CLAMS; Columbus Instruments). Mice were housed individually at 25°C under a 12-h light/12-h dark cycle, with *ad libitum* access to food and water. Locomotor activity was recorded at 1-min intervals from infrared beam breaks along the x-, y-, and z-axes. For hyperinsulinemic– euglycemic clamp studies, mice were surgically implanted with catheters in the carotid artery and jugular vein 7 days before the procedure. Following a 5-h fast, basal glucose turnover was assessed by continuously infusing [3-³H]glucose at 0.05 µCi min⁻¹ for 2 h. A 2-h hyperinsulinemic–euglycemic clamp was subsequently performed with continuous insulin infusion at 4 mU kg⁻¹ min⁻¹. Blood glucose was maintained at approximately 6.7 mM by a variable-rate infusion of glucose supplemented with [3-³H]glucose to maintain stable tracer-specific activity. At the end of the clamp, mice received a bolus of [¹⁴C]2-deoxyglucose ([¹⁴C]2-DG). Tissue-specific glucose uptake was calculated from the accumulation of [¹⁴C]2-DG-6-phosphate in each tissue. Hepatic glucose production was calculated by subtracting the exogenous glucose infusion rate from the total glucose rate of appearance. HOMA2-IR was calculated from fasting glucose and plasma insulin concentrations using the online HOMA2 Calculator. The Adipo-IR index was calculated by multiplying the fasting serum NEFA concentration (nmol l⁻¹) by the fasting serum insulin concentration (ng ml⁻¹).

### Cell Culture

3T3-L1 preadipocytes were maintained in DMEM supplemented with calf serum (10%). To induce differentiation, confluent cells were cultured from days 0–2 in DMEM containing fetal bovine serum (FBS; 10%), IBMX (250 μM), insulin (5 μg mL⁻¹), and dexamethasone (1 μM). From days 2–4, cells were cultured in DMEM containing FBS (10%) and insulin (5 μg mL⁻¹). Beginning on day 4, cells were maintained in DMEM containing FBS (10%) alone^8, 83^. Immortalized murine brown preadipocytes were cultured in DMEM supplemented with FBS (10%) until confluent. Adipogenic differentiation was induced for 2 days in DMEM containing FBS (10%), insulin (850 nM), T3 (2 nM), IBMX (0.5 mM), rosiglitazone (1 μM), and dexamethasone (1 μM). Cells were then maintained for 6 days in DMEM containing FBS (10%), rosiglitazone (1 μM), and T3 (2 nM), with medium changes every 2 days. Human SGBS preadipocytes were cultured on collagen-coated plates in DMEM/F12 containing penicillin– streptomycin (1%), biotin (33 μM), pantothenate (17 μM), and FBS (10%). Two days after confluence, differentiation was induced for 4 days in serum-free medium containing transferrin (0.01 mg mL⁻¹), insulin (20 nM), cortisol (100 nM), T3 (0.2 nM), dexamethasone (25 nM), IBMX (250 μM), and rosiglitazone (2 μM). Cells were then maintained for 4 days in medium containing transferrin (0.01 mg mL⁻¹), insulin (20 nM), cortisol (100 nM), and T3 (0.2 nM) and were considered fully differentiated on day 8. Lentiviral overexpression and CRISPR–Cas9-mediated gene disruption were performed on or after day 8 to minimize potential effects on adipogenesis.

### Insulin and reNRN1 signaling assay

For *in vivo* insulin-signaling studies, mice were fasted overnight and injected intraperitoneally with insulin at 10 U kg⁻¹. Tissues were harvested 10 min after injection. For *ex vivo* signaling studies, freshly dissected tissues were collected in PBS, transferred to DMEM containing 10% FBS, and equilibrated for 3 h. Tissue explants were then treated with recombinant NRN1 (300 ng/mL) for 10 min before collection for downstream analyses.

### ChIP assays

Tissue samples were cross-linked with 1% formaldehyde for 10 min at room temperature. Chromatin was sonicated using an S220 Ultrasonicator (Covaris) to generate DNA fragments of approximately 200–500 bp. An aliquot of each cleared lysate was reserved as input, and the remaining lysate was incubated overnight at 4°C with an antibody against PPARγ or control IgG. Prewashed Dynabeads Protein G (20 µL per immunoprecipitation) were then added, and the samples were rotated for 1 h at 4°C. The beads were sequentially washed with low-salt RIPA buffer (20 mM Tris–HCl, pH 8.0, 1 mM EDTA, 1% Triton X-100, 0.1% SDS, 140 mM NaCl, and 0.1% sodium deoxycholate), high-salt RIPA buffer (20 mM Tris–HCl, pH 8.0, 1 mM EDTA, 1% Triton X-100, 0.1% SDS, 500 mM NaCl, and 0.1% sodium deoxycholate), LiCl buffer (250 mM LiCl, 0.5% NP-40, 0.5% sodium deoxycholate, 1 mM EDTA, and 10 mM Tris–HCl, pH 8.0), and TE buffer (10 mM Tris–HCl, pH 8.0, and 1 mM EDTA). The beads were subsequently incubated for at least 4 h at 65°C in digestion buffer containing 50 mM Tris–HCl, pH 8.0, 1 mM EDTA, 100 mM NaCl, 0.5% SDS, and proteinase K to reverse the cross-links. DNA was recovered by phenol–chloroform extraction, and enrichment was quantified by qPCR using the primers listed in Supplementary Table 1.

### Lipolysis assays

For *in vivo* lipolysis assays, mice were fasted for 4 h and injected intraperitoneally with isoproterenol (10 mg kg⁻¹ body weight). Blood was collected from the tail vein immediately before injection and at 15, 30, and 60 min afterward. Serum glycerol and nonesterified fatty acid (NEFA) concentrations were measured using the Free Glycerol Determination Kit (Sigma-Aldrich) and the HR Series NEFA-HR Kit (FUJIFILM), respectively. *Ex vivo* lipolysis was assessed as previously described, with minor modifications^84^. Freshly excised adipose tissue was cut into small pieces (150–300 mg) and incubated in DMEM containing 10% FBS for 3 h. The medium was then replaced with DMEM containing either vehicle or reNRN1 (300 ng ml⁻¹), and the tissue was incubated for 24 h. Next, the tissue was transferred to prewarmed Krebs– HEPES Ringer buffer containing 2% fatty acid-free BSA (KRBH–BSA) and equilibrated for 1 h. A 100-µl aliquot of medium was collected as the baseline sample. The tissue was then incubated in KRBH–BSA containing 10 µM isoproterenol, and 100-µl aliquots were collected at 30, 60, and 120 min. All aliquots were heated at 65 °C for 10 min to terminate enzymatic activity. Glycerol and NEFA concentrations in the medium were measured using the kits described above.

### Co-immunoprecipitation

Cells were washed with ice-cold PBS and lysed in 1 mL of buffer containing 50 mM Tris–HCl, pH 7.5, 150 mM NaCl, 1% Triton X-100, and a protease inhibitor cocktail for 30 min on ice with continuous rotation. Lysates were clarified by centrifugation at 15,000 rpm for 10 min at 4°C, and the resulting supernatants were incubated with anti-FLAG antibody–conjugated beads (Sigma-Aldrich) for 2 h at 4°C. The beads were washed four times with lysis buffer, and bound proteins were eluted by boiling in SDS sample buffer. Eluted proteins were separated by SDS– PAGE and analyzed by immunoblotting using standard procedures.

### Immunofluorescence

Cells grown on glass coverslips were fixed with 4% paraformaldehyde. As indicated, cells were either permeabilized with PBS containing 0.1% Triton X-100 for 10 min or left nonpermeabilized. Cells were incubated for 18 h at 4°C with the indicated primary antibodies diluted in PBS containing 10% (v/v) FBS. After washing, the cells were incubated for 1 h with FITC-conjugated anti-rabbit IgG or Cy3-conjugated anti-mouse IgG secondary antibodies, as appropriate. Fluorescence images were acquired using a Zeiss LSM 710 confocal microscope (Carl Zeiss).

### 2-DG assay

Cells, tissue explants, and mature adipocytes were incubated in serum-free DMEM for 4–6 h. Samples were washed three times with Krebs–Ringer HEPES (KRH) buffer containing 12 mM HEPES, pH 7.4, 121 mM NaCl, 5 mM KCl, 0.33 mM CaCl₂, and 1.2 mM MgSO₄ and then incubated for 20 min in KRH buffer with or without 100 nM insulin. [2,6-³H]2-Deoxy-D-glucose was subsequently added at 0.33 μCi mL⁻¹, and uptake was allowed to proceed for 10 min. Uptake was terminated by three rapid washes with ice-cold KRH buffer containing 200 mM glucose and 10 μM cytochalasin B. Samples were solubilized in 0.1% SDS for 30 min, and radioactivity was measured by liquid scintillation counting.

### PI-PLC treatment

HEK293T cells were transfected with an NRN1 expression plasmid using Lipofectamine 2000 according to the manufacturer’s instructions. After 24 h, the cells were washed with PBS and incubated in serum-free medium containing vehicle or 0.1 U/mL PI-PLC for 2 or 16 h. Following treatment, conditioned media were concentrated, and cell membrane fractions were isolated for subsequent analyses^85^.

### Membrane fractionation

Cells were washed with PBS, and membrane fractions were isolated using the Cell Fractionation Kit (Cell Signaling Technology, catalog no. 9038) according to the manufacturer’s instructions.

### Phopho-kinase array profiling

Freshly isolated mouse adipose tissue was incubated in DMEM containing 10% FBS for 3 h and then treated with recombinant NRN1 (reNRN1; 300 ng/mL) for 10 min. The tissue was washed with PBS and subjected to phospho-kinase array profiling using the Proteome Profiler Phospho-Kinase Array Kit (ARY003C; R&D Systems) according to the manufacturer’s instructions.

### Immunoblotting

Cell and tissue samples were lysed in RIPA buffer containing 50 mM Tris–HCl, pH 7.5, 150 mM NaCl, 1% Triton X-100, 0.1% SDS, and cOmplete Mini EDTA-free Protease Inhibitor Cocktail (cat. no. 11836170001; Roche) for 30 min on ice with continuous rotation. Protein concentrations were determined using the Bradford assay. For each sample, 40–60 μg of protein was separated by SDS–PAGE and transferred to a polyvinylidene difluoride or nitrocellulose membrane. Membranes were blocked with 5% nonfat dry milk in Tris-buffered saline containing 0.25% Tween 20 (TBST) and incubated overnight with the indicated primary antibodies. After washing, membranes were incubated with the appropriate secondary antibodies for 1 h at room temperature and washed with TBST for 30 min. Chemiluminescent signals were developed using ECL reagent (NEL104001EA; PerkinElmer), captured using an iBright CL1500 Imaging System (Invitrogen), and quantified using ImageJ.

### Reagents

Insulin, dexamethasone, IBMX, rosiglitazone, BGJ398, and isoproterenol were purchased from Sigma-Aldrich. Recombinant NRN1 proteins were purchased from Novus Biologicals (NBP2-35073). The following antibodies were used: HSP90 (Cell Signaling Technology, 4874), GAPDH (Cell Signaling Technology, 2118), mouse IgG1 isotype control (clone G3A1; Cell Signaling Technology, 5415), FLAG (Sigma-Aldrich, F3165), GFP (Abcam, ab290), PPARγ (Cell Signaling Technology, 2443), NRN1 (Sigma-Aldrich, PRS4101), pan-cadherin (Cell Signaling Technology, 4068), PGC-1α (R&D Systems, 474915), phospho-AKT (Cell Signaling Technology, 9271), AKT (Cell Signaling Technology, 9272), phospho-HSL (Cell Signaling Technology, 4139), UCP1 (Cell Signaling Technology, 72298), FGFR1 (Cell Signaling Technology, 9740), HA (Cell Signaling Technology, 3724), ERK1/2 (Cell Signaling Technology, 9102), His tag (Cell Signaling Technology, 12698), phospho-p38 (Santa Cruz Biotechnology, sc-166182), phospho-IRS1 (Invitrogen, 44-816G), and phospho-insulin receptor (IR) (Invitrogen, 44-800G).

### RNA extraction and qPCR

Total RNA was extracted from tissues using TRIzol reagent according to the manufacturer’s instructions. Complementary DNA (cDNA) was synthesized from 1 μg of total RNA using the High-Capacity cDNA Reverse Transcription Kit (Applied Biosystems). Quantitative PCR (qPCR) was performed using 2× SYBR Green qPCR Master Mix (Thermo Fisher Scientific) on a QuantStudio 5 Real-Time PCR System (Applied Biosystems). Relative mRNA expression was normalized to cyclophilin B and calculated using the 2^-ΔΔCt^ method. Primer sequences are provided in Supplementary Table 1.

### Plasmids

Lentiviral expression constructs encoding NRN1, FGFR1, KLB, NRN1-ΔGPI, and NRN1^G116E^ were generated by cloning the corresponding cDNAs into the pCDH-CMV-MCS-EF1 vector using XbaI/BamHI for NRN1, NRN1-ΔGPI, and NRN1^G116<E^; KpnI/NotI for NRN1-TM; XbaI/NotI for FGFR1; and EcoRI/NotI for KLB. Single-guide RNAs (sgRNAs) targeting the genes of interest were cloned into the lentiCRISPR v2 vector. Briefly, lentiCRISPR v2 was digested with BsmBI and dephosphorylated, and oligonucleotides encoding the sgRNAs were phosphorylated, annealed, and ligated into the digested vector. Hairpin and sgRNA sequences are provided in **Supplementary Table 1**. Lentiviral particles were produced using the packaging plasmids psPAX2 and pMD2.G (Addgene).

### Insulin ELISA

For insulin measurements, mice were fasted for 6 h before blood collection, and plasma insulin concentrations were measured using an Ultra Sensitive Mouse Insulin ELISA Kit (Crystal Chem, 90080). Assays were performed according to the manufacturer’s instructions, and endpoint absorbance was measured using a SpectraMax i3 plate reader (Molecular Devices).

### Cell-surface crosslinking with DTSSP and His-tag pulldown

3T3-L1 adipocytes were cultured in type I collagen-coated 10-cm dishes and serum-starved for 4 h. Cells were chilled on ice for 10 min and incubated with 10 nM His-tagged NRN1 or the indicated control protein for 20 min. After two washes with 15 mL of ice-cold PBS, cell-surface proteins were cross-linked with 2.5 mM 3,3′-dithiobis(sulfosuccinimidyl propionate) (DTSSP) for 30 min at room temperature. Cross-linking was quenched with 20 mM Tris–HCl, pH 7.5. Cells were lysed in 1 mL of RIPA buffer containing protease and phosphatase inhibitor cocktails. Lysates were passed ten times through a 22-gauge needle, rotated for 20 min at 4°C, and clarified by centrifugation at 15,000 × *g* for 10 min. Imidazole was added to a final concentration of 10 mM, and His-tagged protein complexes were captured using 100 μL of Ni–NTA agarose for 1 h at 4°C. The beads were washed once with RIPA buffer containing 10 mM imidazole and three times with RIPA buffer containing 30 mM imidazole. Bound proteins were eluted with 800 μL of RIPA buffer containing 250 mM imidazole for 20 min at 4°C. Eluted proteins were precipitated on ice for 1 h by adding 100 μL of 0.2% sodium deoxycholate and 100 μL of 10% trichloroacetic acid. Protein pellets were collected by centrifugation at 12,000 × *g* for 10 min at 4°C, washed twice with ice-cold acetone, dried at 37°C, and resuspended in PBS and NuPAGE LDS sample buffer containing 5 mM DTT. Samples were heated at 65°C for 20 min and alkylated with 14 mM iodoacetamide for 45 min in the

### Transcriptomics studies

Total RNA was extracted using the RNAqueous-Micro Kit (cat. no. AM1931), and RNA integrity was assessed using an Agilent 2100 Bioanalyzer. Poly(A)-selected libraries were prepared using the Poly(A) RNA Selection Kit v1.5. Sequencing reads were aligned to the UCSC mm39 reference genome using HISAT2^86^, and gene-level counts were generated using HTSeq^87^. Differential-expression analysis was performed using DESeq2^88^. Differentially expressed genes (DEGs) were considered by a false discovery rate (FDR) below 0.05. To identify pathways showing opposite regulation between the knockout and transgenic models, DEGs were classified as upregulated or downregulated in each model and intersected according to their direction of change. Over-representation analysis (ORA) of these gene sets was performed using GSEApy^89^ with the KEGG 2019 Mouse gene sets.

## Statistical analysis

Data are presented as mean ± SEM. Statistical comparisons were made using unpaired two-tailed Student’s t-test for two-group comparisons two-way ANOVA with appropriate post-hoc correction for multi-group comparisons. All statistical analyses were performed in GraphPad Prism 9 or R version 4.2.

## Statistical analysis

Data are presented as mean ± SEM. Statistical comparisons were made using unpaired two-tailed Student’s t-test for two-group comparisons or one-way/two-way ANOVA with appropriate post-hoc correction for multi-group comparisons. Correlation analyses used Pearson’s coefficient. A p-value <0.05 was considered statistically significant. All statistical analyses were performed in GraphPad Prism 9 or R version 4.2.

## Data Availability

Raw RNA-sequencing data have been deposited in the Gene Expression Omnibus under accession number GSE346412.

## Acknowledgements

We thank Drs. Evan D. Rosen and Hei Sook Sul for reviewing the manuscript.

## Ethics Declaration

The authors declare no competing interests.

