## Supplementary Figures and table for "Neuritin-1 is an adipokine that sustains insulin sensitivity through adipocyte FGFR1–PPARγ signaling"

Supplementary Figure 1

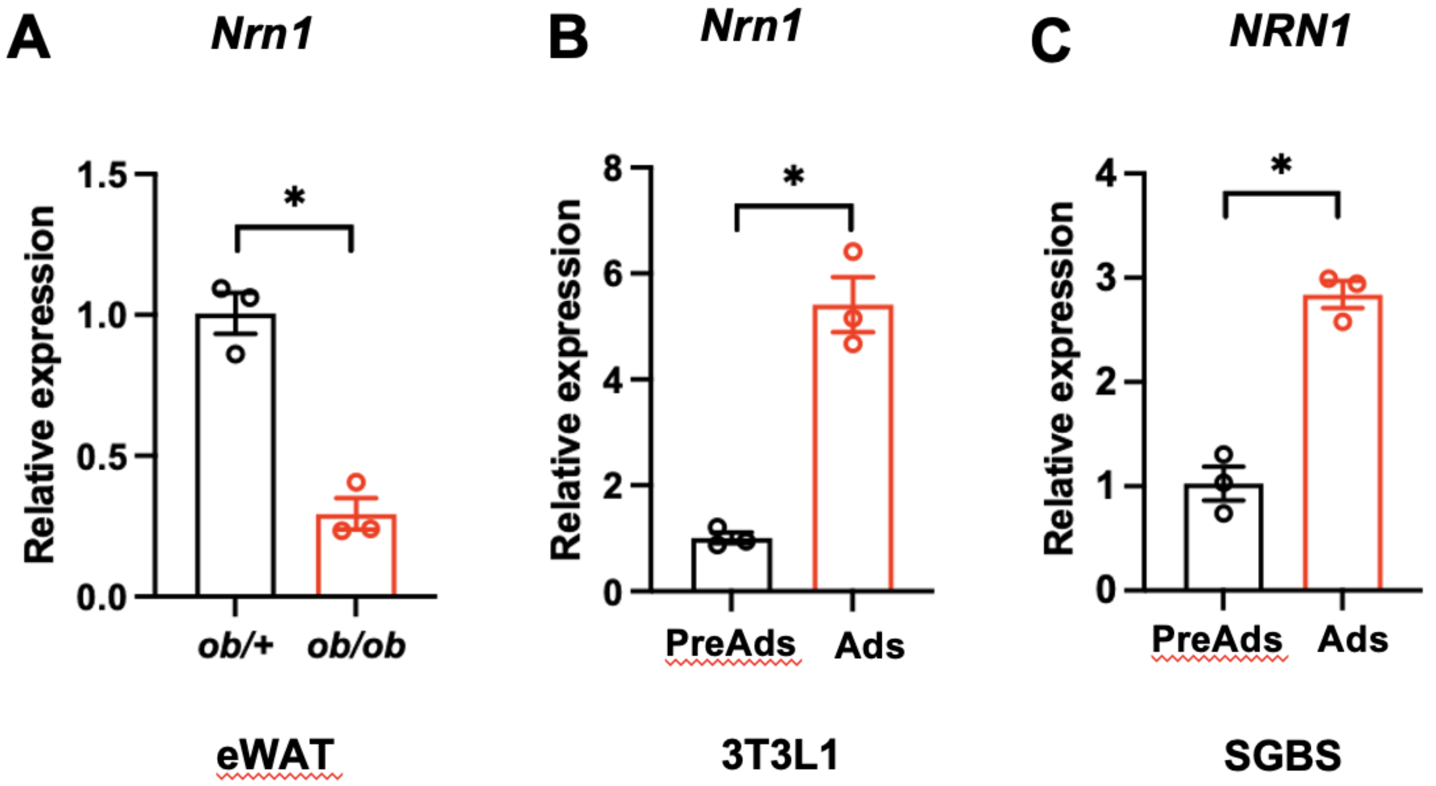

**Supplementary Figure 2**

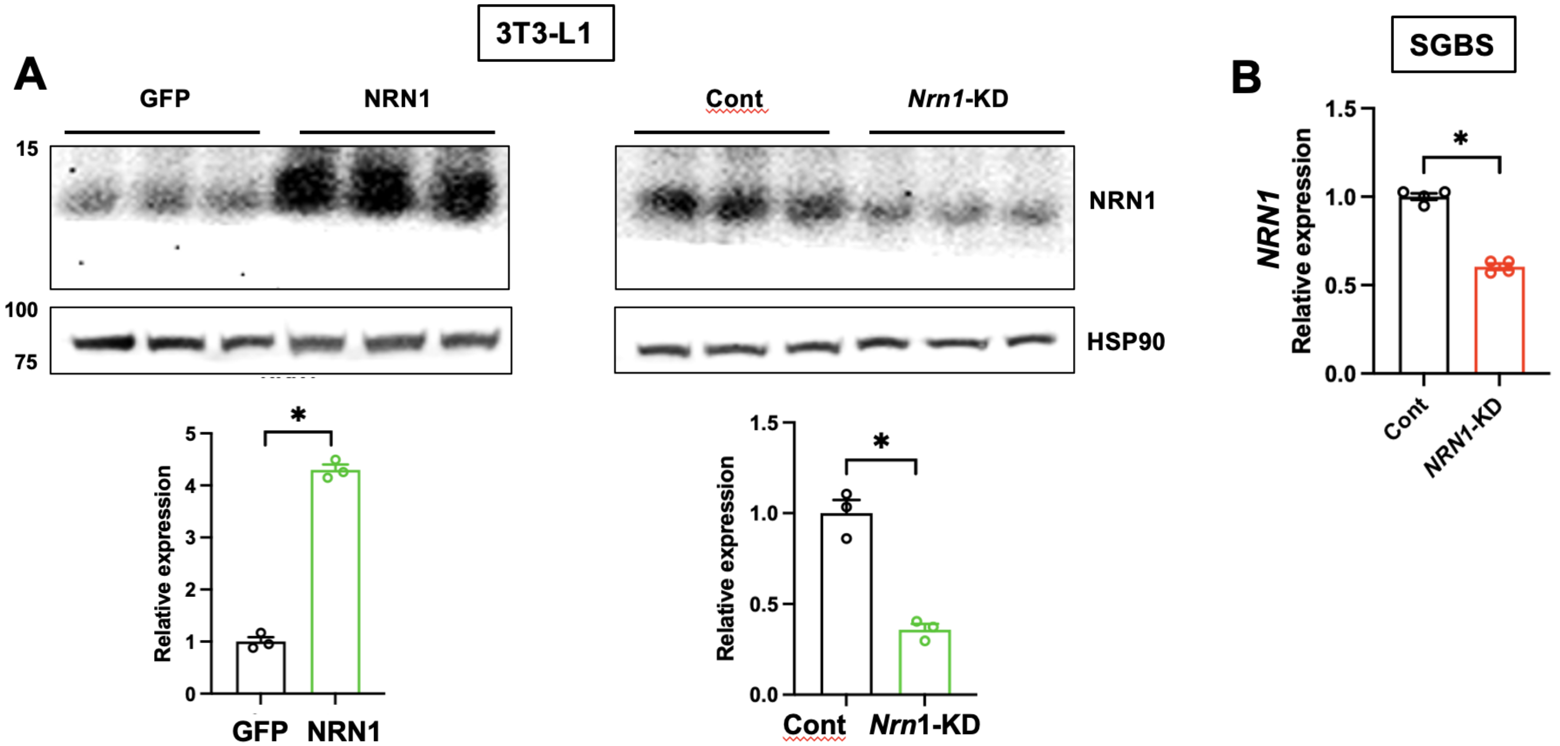

Supplementary Figure 3

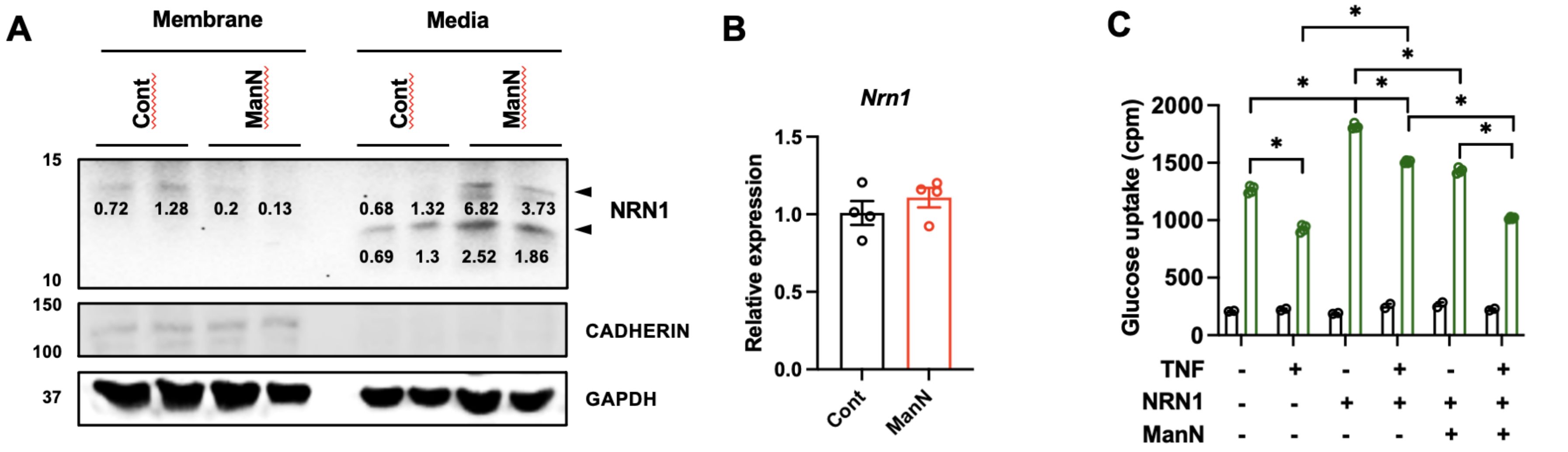

Supplementary Figure 4

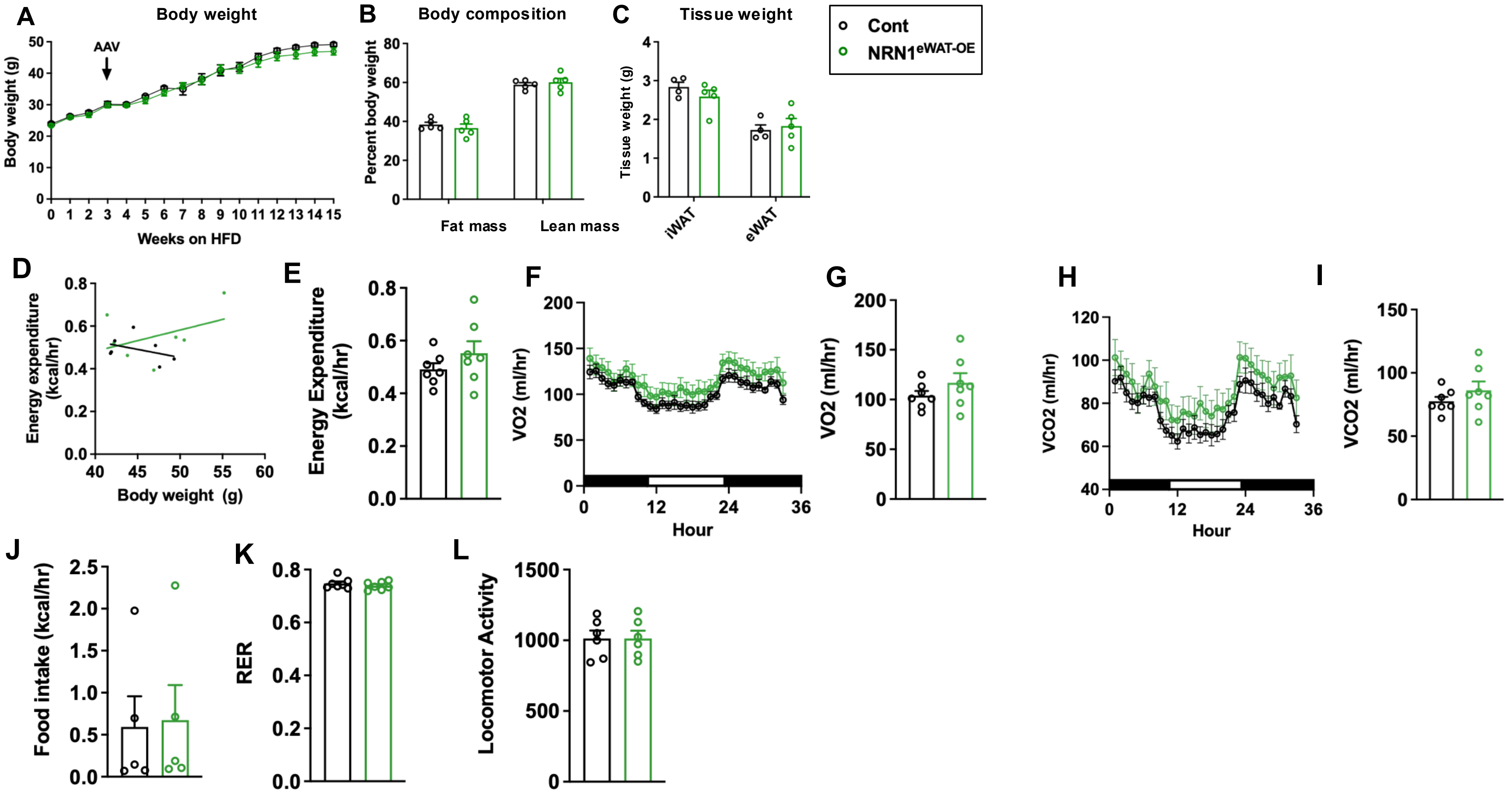

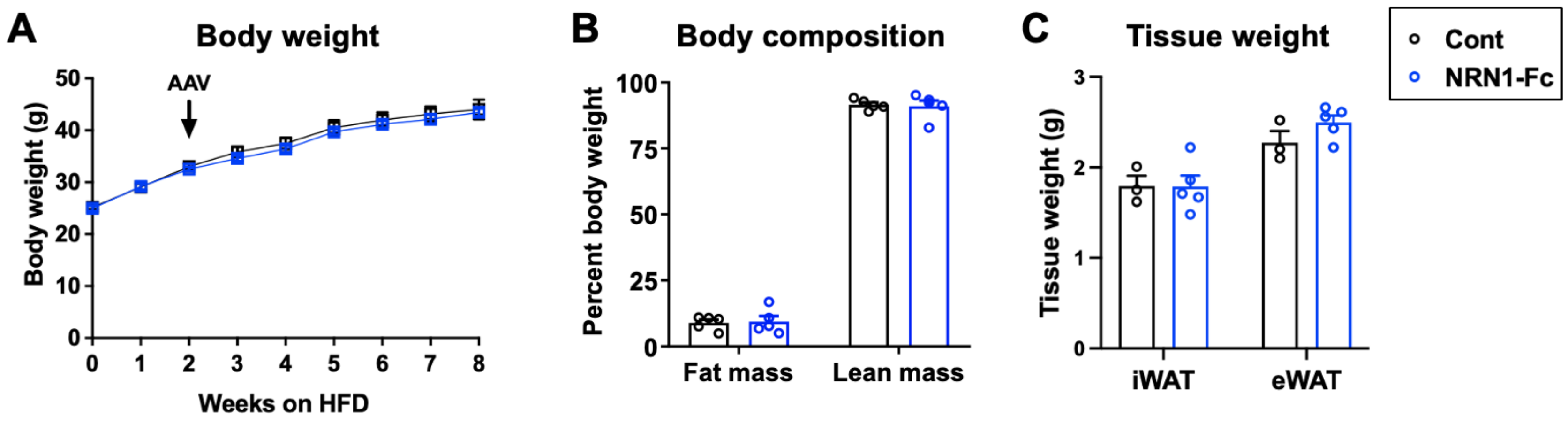
A
Body weight
AAV
Body weight (g)
50
40
30
20
10
0
0
1
2
3
4
5
6
7
8
Weeks on HFD
B
Body composition
Percent body weight
100
75
50
25
0
Fat mass
Lean mass
C
Tissue weight
Tissue weight (g)
3
2
1
0
iWAT
eWAT
Cont
NRN1-Fc

### Supplementary Figure 6

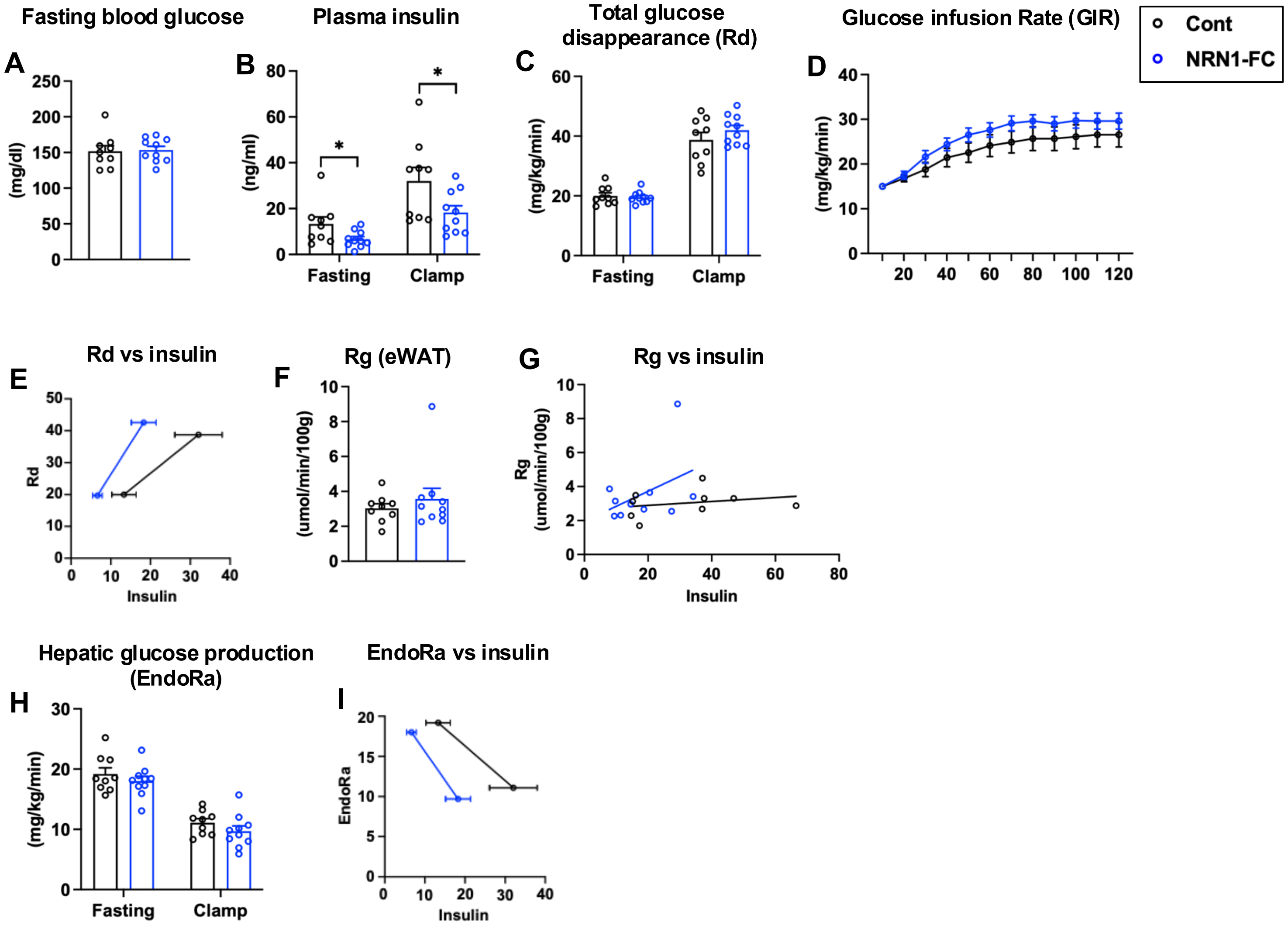

### Supplementary Figure 7

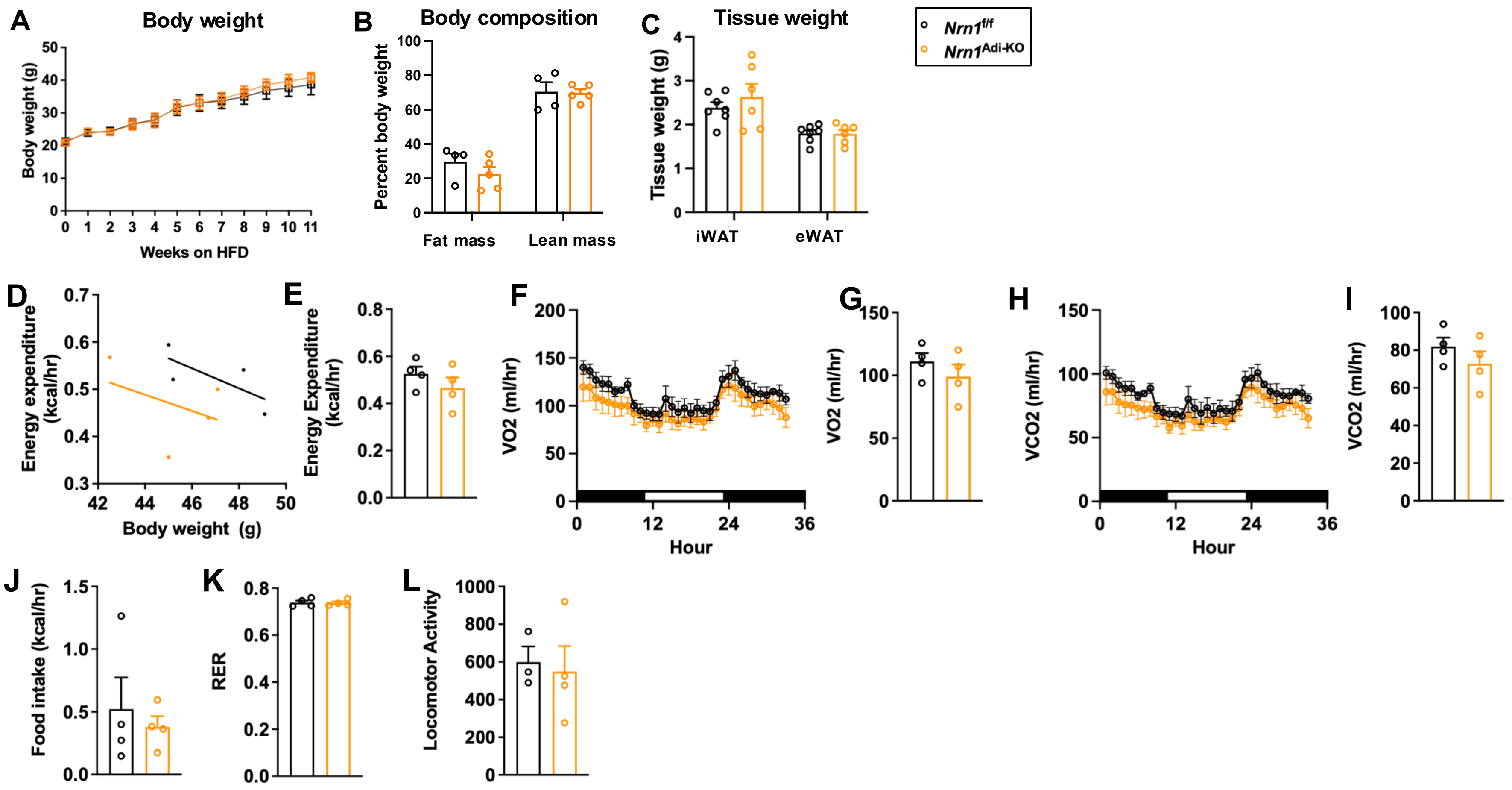

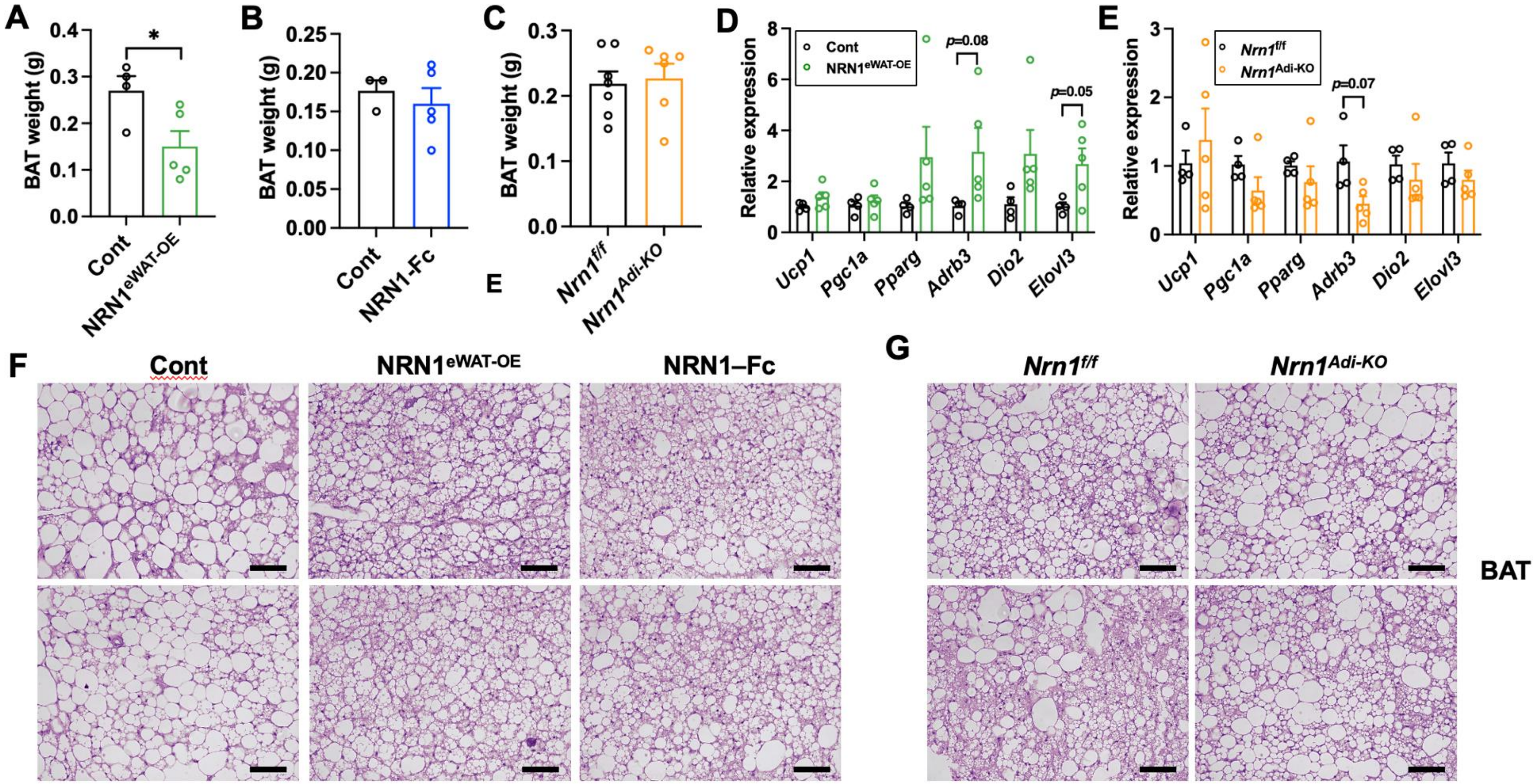

A
B
C
D
E
F
G
BAT weight (g)
Cont
NRN1eWAT-OE
NRN1-Fc
Nrn1f/f
Nrn1Adi-KO
Relative expression
p=0.08
p=0.05
p=0.07
Ucp1
Pgc1a
Pparg
Adrb3
Dio2
Elovl3
NRN1–Fc
BAT

### Supplementary Figure 9

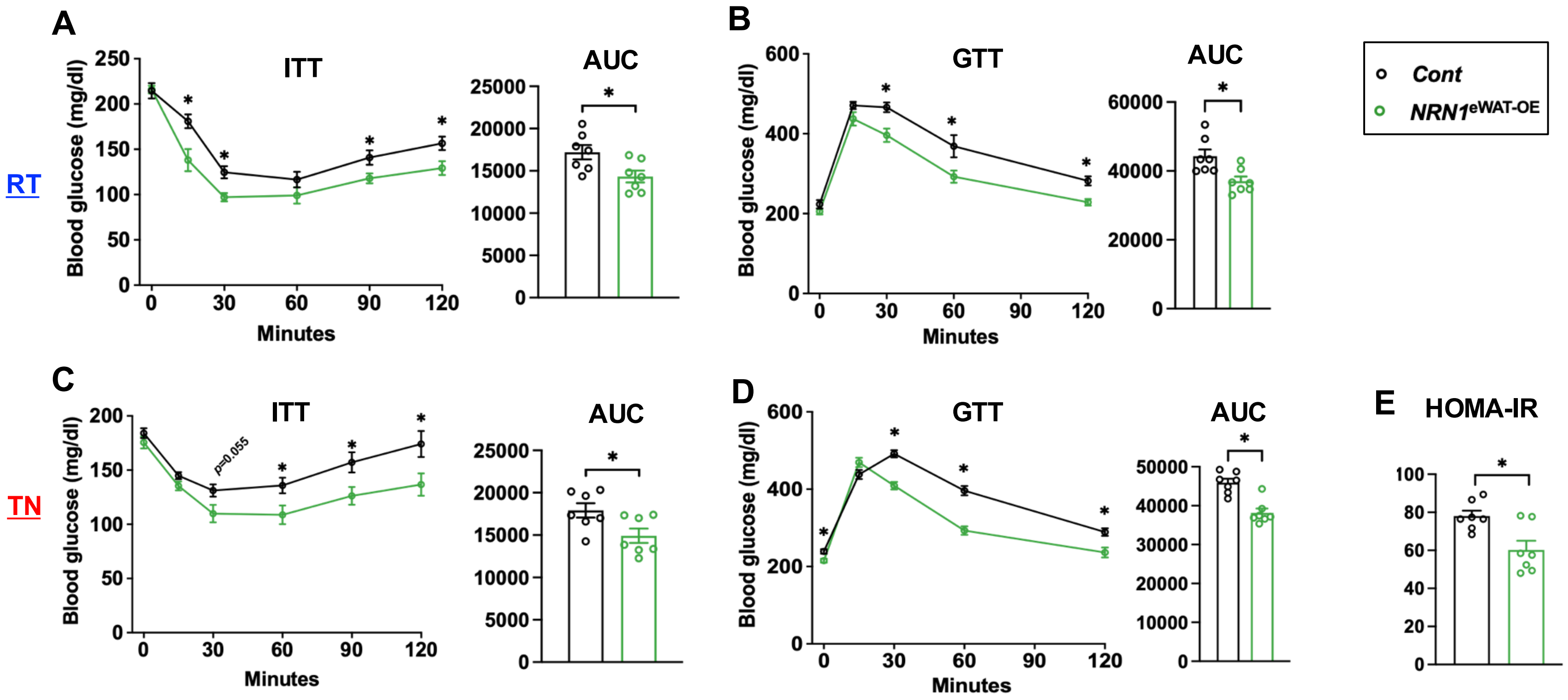

### Supplementary Figure 10

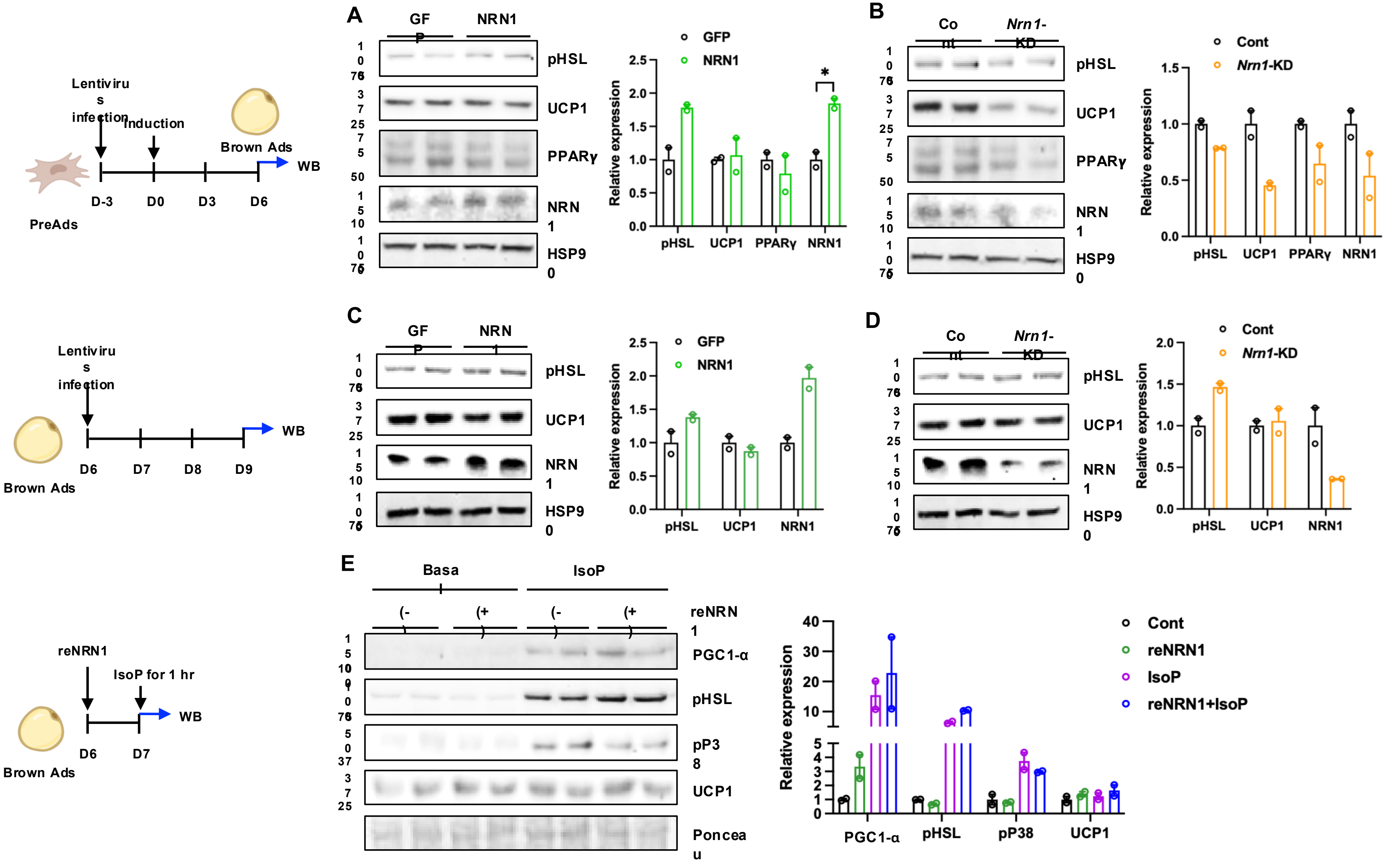

**Supplementary Figure 11**

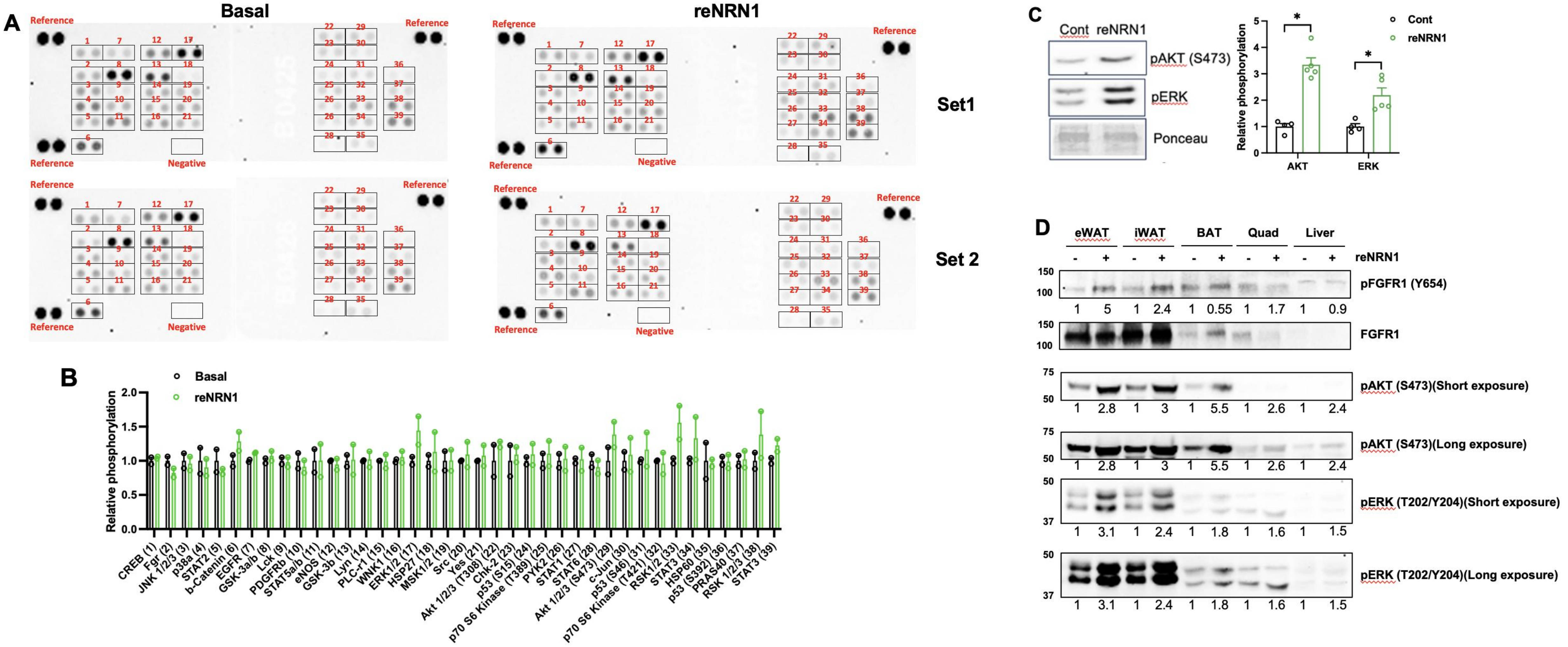

**Supplementary Figure 12**

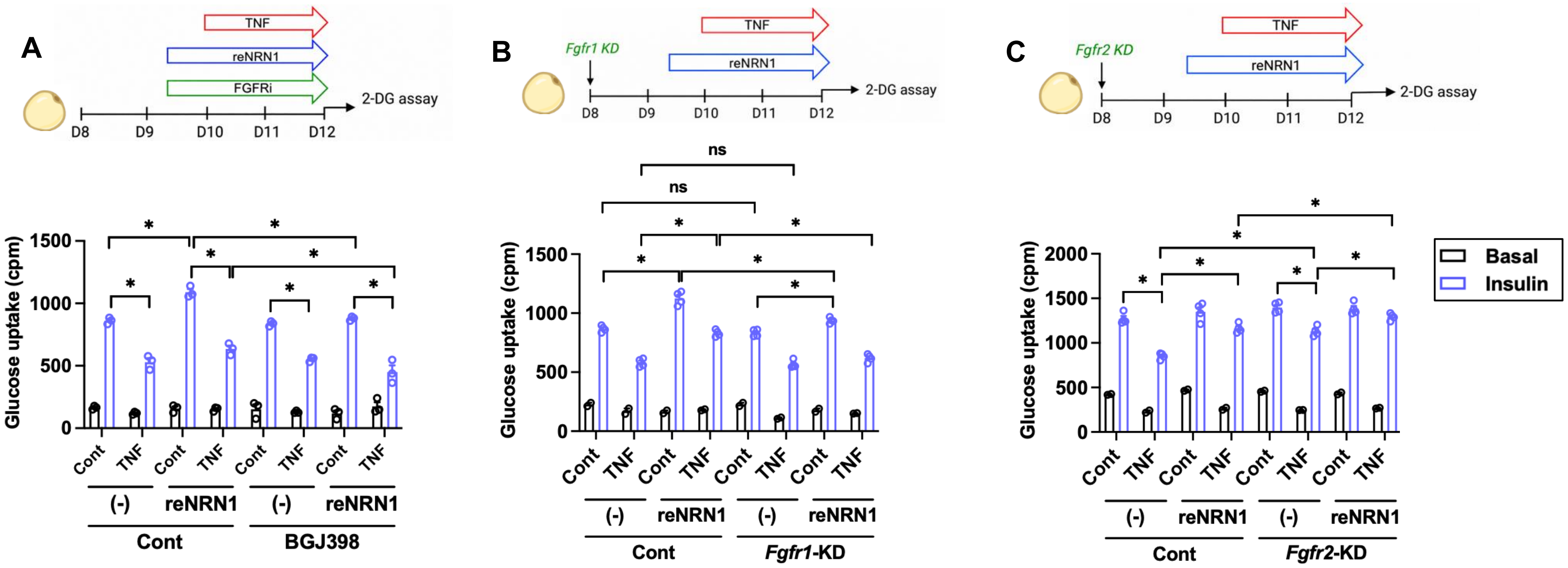

**Supplementary Figure 13**

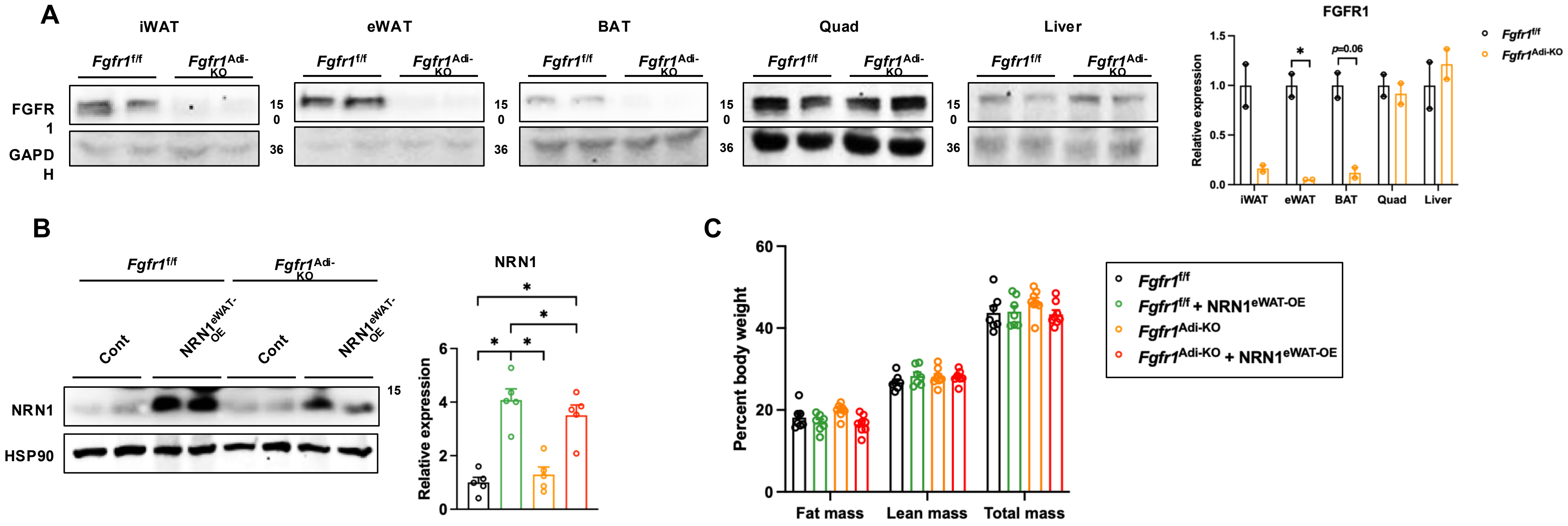

**Supplementary Figure 14**

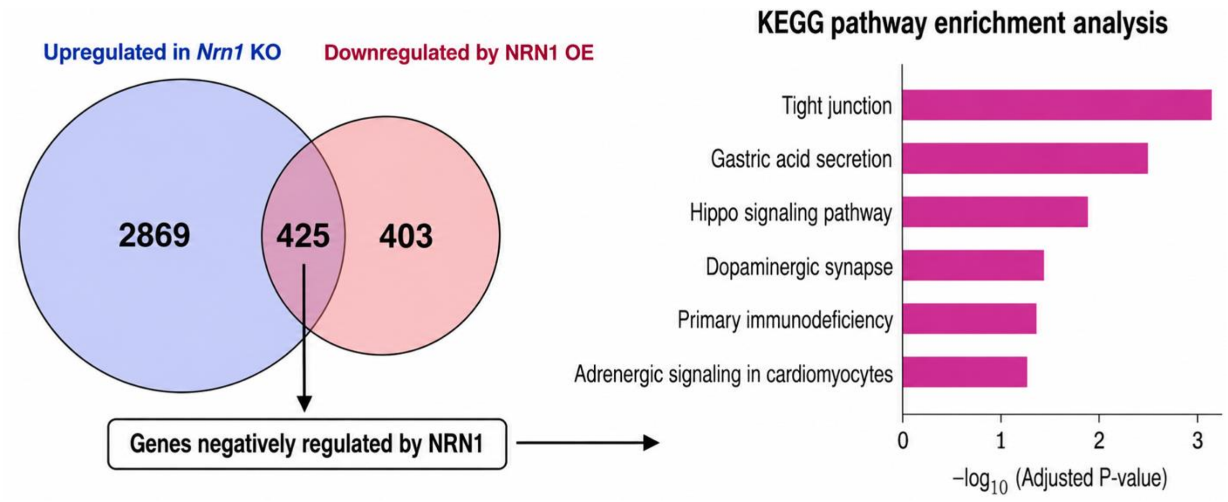

**Supplementary Table 1. Oligonucleotide sequences used in this study.**

| | | |
|---|---|---|
| qPCR | Mouse Nrn1_F | GCGGTGCAAATAGCTTACCTG |
| | Mouse Nrn1_R | CGGTCTTGATGTTCGTCTTGTC |
| | Human NRN1_F | GCGGTGCAAATAGCGTATCTG |
| | Human NRN1_R | CGGTCTTGATGTTCGTCTTGTC |
| | Il6_F | CTCTGGGAAATCGTGGAAAT |
| | Il6_R | CCAGTTTGGTAGCATCCATC |
| | Ccl2_F | GGCCTGCTGTTCACAGTTGC |
| | Ccl2_R | CCTGCTGCTGGTGATCCTCTT |
| | Tnf_F | ATGAGAAGTTCCCAAATGGC |
| | Tnf_R | CTCCACTTGGTGGTTTGCTA |
| | Il1b_F | GCAACTGTTCCTGAACTCAACT |
| | Il1b_R | ATCTTTTGGGGTCCGTCAACT |
| | Pparg_F | CAAGAATACCAAAGTGCGATCAA |
| | Pparg_R | GAGCTGGGTCTTTTCAGAATAATAAG |
| | Adipoq_F | TGTTCCTCTTAATCCTGCCCA |
| | Adipoq_R | CCAACCTGCACAAGTTCCCTT |
| | Fabp4_F | AAGGTGAAGAGCATCATAACCCT |
| | Fabp4_R | TCACGCCTTTCATAACACATTCC |
| | Lipe_F | CCGCTGACTTCCTGCAAGAG |
| | Lipe_R | CTGGGTCTATGGCGAATCGG |
| | Fgfr1_F | GACCACAGAATTGGAGGCTA |
| | Fgfr1_R | ATGAACTCCACGTTGCTACC |
| | Fgf1_F | CCCTGACCGAGAGGTTCAAC |
| | Fgf1_R | GTCCCTTGTCCCATCCACG |
| | Fgf21_F | AGATCAGGGAGGATGGAACA |
| | Fgf21_R | TCAAAGTGAGGCGATCCATA |
| | Glut4_F | TCATTGTCGGCATGGGTTT |
| | Glut4_R | CGGCAAATAGAAGGAAGACGT |
| | Ucp1_F | CACCTTCCCGCTGGACACT |
| | Ucp1_R | CCCTAGGACACCTTTATACCTAATGG |
| | Pgc1a_F | AGCCGTGACCACTGACAACGAG |
| | Pgc1a_R | GCTGCATGGTTCTGAGTGCTAAG |
| | Adrb3_F | TGCGCACCTTAGGTCTCATTAT |
| | Adrb3_R | AAGGCGGAGTTGGCATAGC |
| | Dio2_F | CAGTGTGGTGCACGTCTCCAATC |
| | Dio2_R | TGAACCAAAGTTGACCACCAG |
| | Elovl3_F | TCCGCGTTCTCATGTAGGTCT |
| | Elovl3_R | GGACCTGATGCAACCCTATGA |
| | Pck1_F | AAGTGCCTGCACTCTGTGG |
| | Pck1_R | CAGGCCCAGTTGTTGACC |
| | G6pc_F | TCCCTGTCACCTGTGAG |
| | G6pc_R | TTACAAAGACTTCTTGTG |
| | Cyclophilin_F | GGTGGAGAGCACCAAGACAGA |
| | Cyclophilin_R | GCCGGAAGTCGACAATGATG |
| | Plin1_F | GATCGGCTCTGAACTGAAGG |
| | Plin1_R | CTTCTCGATGCTTCCCAGAG |

| | | |
|---|---|---|
| ChIP/MeDIP | Nrn1_#1_F | GAGAAGGCAAGGACGTTGAG |
| | Nrn1_#1_R | AAGCTCTTCCCCCATACTCC |
| | Nrn1_#2_F | AGCGTCGCGAATAGTGACAT |
| | Nrn1_#2_R | CTAGAGCGCGTGAGCACATA |
| | Ins_F | CTTCAGCCCAGTTGACCAAT |
| | Ins_F | AGGGAGGAGGAAAGCAGAAC |
| ChIP | Pparg_F | GGACTCCATTACATCCTTAAGTAG |
| | Pparg_R | CGAGAAGTTGGCAATTACATC |
| gRNA | Human NRN1 | GTCCAGGCCCTGCGGGTAGTTGG |
| | Mouse Fgfr2 | AGATGGGATTACCGTCCACGTGG |
| shRNA | Mouse Nrn1 | AAGGGCTTTTCAGACTGTTTG |
| | Mouse Fgfr1 | CCTGGAGCATCATAATGGATT |
